# Global Diversity in Mammalian Life Histories: Environmental Realms and Evolutionary Adaptations

**DOI:** 10.1101/2023.06.29.546851

**Authors:** E. Beccari, P. Capdevila Lanzaco, R. Salguero-Gómez, C. Pérez Carmona

**Author notes:** Author for correspondence: Eleonora Beccari.

## Abstract

Mammalian life history strategies can be characterized by a few axes of variation, which conform a space where species are positioned according to which life history strategies are favoured in the environment they exploit. Yet, we still lack global descriptions of the diversity of realized mammalian life history and of how this diversity is shaped by the environment. We used six life history traits to build a global life history space and explored how major environmental realms (land, air, water) influence mammalian life history strategies. We demonstrate that realms are tightly linked to distinct life history strategies. Predominantly, aquatic and aerial species adhere to slower life history strategies, while terrestrial species tend to exhibit faster life histories. Highly encephalized terrestrial species are a notable exception to these patterns. In addition, species transitioning between the terrestrial and aquatic realms, such as seals, show intermediate life history strategies. Further, different mode-of-life may play a significant role in allowing to expand the set of strategies exploitable in the terrestrial realm. Our results provide compelling evidence of the link between environmental realms and the diversity of life history strategies among mammals.

**Statement of authorship:** P.C.L., R.S-G., and C.P.C. conceived the core ideas behind this paper, all authors provided fundamental inputs for its final development. E.B collected all data and performed the literature search needed to sort species in environmental realms. E.B. analysed the data with fundamental inputs from C.P.C., P.C.L, and R.G-S. All authors contributed to interpretation of the results. E.B. and C.P.C. led the writing of the manuscript which was edited by all authors.

## Introduction

The diversity of mammalian adaptations, ranging from the fast and semelparous life histories of antechinus (Braithwaite & Crowther 2008) to the slow and long lives of bowhead whales (Perrin *et al*. 2009), have allowed mammals to occupy all land (Burgin *et al*. 2018; Grossnickle *et al*. 2019), exploit air (Babich Morrow *et al*. 2021; Barclay 1994; Maina 2000), and even return to the sea (Davis 2019). Despite this wide ecological radiation (Grossnickle *et al*. 2019), evolutionary history and energetic constraints limit the set of life history strategies (*i.e.,* combinations of life history traits) that are viable in nature (Grime & Pierce 2012; Southwood 1988; Stearns 1992). These limits restrict mammalian investments in survival, growth, and reproduction to two major dimensions of life history trait variation (Carmona *et al*. 2021b; Gaillard *et al*. 1989; Healy *et al*. 2019; Oli 2004). The first dimension generally encompasses the time scale of life, referred to as the fast-slow continuum (Bielby *et al*. 2007; Capdevila *et al*. 2020; Gaillard *et al*. 1989; Healy *et al*. 2019; Stearns 1992). In contrast, the second dimension depends more on the subset of species and traits considered, but generally reflects the timing and intensity of a species’ reproductive investment (Bielby *et al*. 2007; Capdevila *et al*. 2020; Gaillard *et al*. 1989; Healy *et al*. 2019). While these two life history dimensions are consistently observed across studies with various species and trait subsets (Bielby *et al*. 2007; Capdevila *et al*. 2020; Gaillard *et al*. 1989), we still lack a comprehensive understanding of how mammals are organized within this broad life history space. This gap hampers exploring and quantifying realized mammalian life history diversity, preventing a thorough comprehension of similarities and differences among mammalian species (Carmona *et al*. 2021b; Pinsky *et al*. 2022).

Environmental conditions profoundly influence species’ life history strategies (Grime & Pierce 2012; Southwood 1988). Variable environments, for instance, affect the pace of life of species (Tuljapurkar *et al*. 2009) selecting for delayed life cycles and longer lives compared to constant and stable environments (Grime & Pierce 2012; Tuljapurkar *et al*. 2009). Because life history strategies are selected by similar environmental pressures (Grime & Pierce 2012; Southwood 1988), there is a highly redundant occupation of the life history space regardless of taxonomic affiliation (Carmona *et al*. 2021b; Cooke *et al*. 2019; Cox *et al*. 2021). However, mammal’s widespread distribution across major environmental realms (*i.e*., land, water, air), begs the question of whether mammalian life history strategies are equally represented across these three realms.

Environmental realms are characterized by distinct physical dimensions, structural properties, and environmental requirements (Pinsky *et al*. 2022; Steele 1985; Webb 2012). Consequently, species inhabiting each specific environmental realm are under unique eco-physiological constraints that are not experienced by species from other realms (Pinsky *et al*. 2019; Webb 2012). For instance, the energetically expensive nature of the aquatic and aerial realms (Barclay 1994; Gearty *et al*. 2018; Goldbogen 2018; Guigueno *et al*. 2019; Maina 2000) has led to longer lives in aerial species (Babich Morrow *et al*. 2021; Gaillard *et al*. 1989) or long age at female maturity in the aquatic realm (Davis 2019; Perrin *et al*. 2009). Accordingly, inhabiting different environmental realms may select distinct sets of life history strategies (Capdevila *et al*. 2020; Pinsky *et al*. 2022). Yet, some similarities do exist across realms. For example, the capacities for climbing trees or burrowing on the ground offer partial escape routes from the environmental constrains unique to terrestrial living, selecting for generally longer lifespans compared to other terrestrial species (Bels & Russell 2023; Healy *et al*. 2014, 2019; Mincer & Russo 2020; Withers *et al*. 2016). Similarly, larger brains generally allows species to avoid hazardous situations, selecting for slower lives compared to other terrestrial representatives (Benson-Amram *et al*. 2016; Bertrand *et al*. 2022; Dembitzer *et al*. 2022; González-Lagos *et al*. 2010; Seyfarth & Cheney 2002). It becomes clear how a comprehensive understanding of mammalian adaptations across various environmental realms is crucial for unravelling the complexities of life history strategies. Without this understanding, we will remain ill-equipped to reveal overarching ecological patterns or to understand the driving forces and consequences of ongoing global changes (Pinsky *et al*. 2022).

Here, we use six life history traits representing investments in survival, growth, and reproduction for 3,438 mammalian species (53.7% of currently recognized living mammals, Burgin *et al*. 2018) extracted from the Amniote dataset (Myhrvold *et al*. 2015) to examine the influence of major environmental realms on mammalian life history strategies. In agreement with previous research (Bielby *et al*. 2007; Gaillard *et al*. 1989), we first show that two axes of variation relating to the fast-slow continuum and the reproductive strategies continuum are sufficient to explain a large proportion of mammalian life history strategies (Fig. 1a). Using this plane as a springboard, we proceed to test three hypotheses: (H1) species’ position in the life history space will associate with the environmental realm they occupy, as a consequence the different eco-physiological pressures acting among major realms (Grime & Pierce 2012; Pinsky *et al*. 2022; Tuljapurkar *et al*. 2009; Webb 2012). Specifically, we expect aerial species to exhibit slower life history strategies compared to terrestrial species, and aquatic species to occupy areas of the life history space skewed toward higher iteroparity (Capdevila *et al*. 2020). (H2) Because species transitioning between aquatic and terrestrial environments (*i.e.,* semi-aquatic species) experience a blend of selective pressures, they will occupy intermediate positions in the life history space between these two realms (Hood 2020; Perrin *et al*. 2009). Additionally, (H3) within the terrestrial realm, species with arboreal, semi-arboreal and fossorial mode-of-life will display slower life history strategies than species confined on land.

**Fig. 1.**
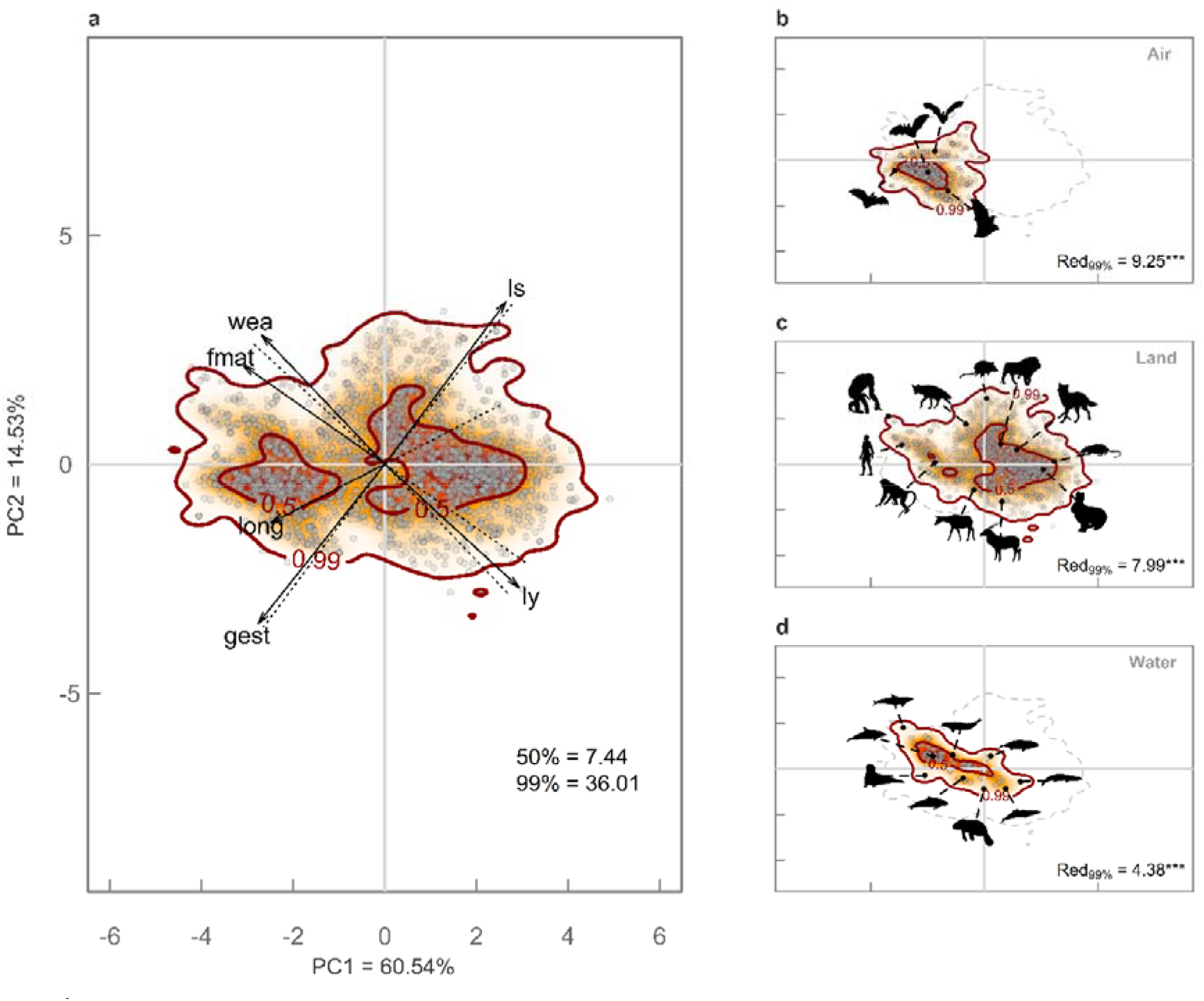
Bimodal global mammal space occupation stems from differences among environmental realms. **a**, Probabilistic species distributions in the space defined by the two first principal components (PC1 and PC2) of PCA considering life history traits for all mammals. Arrows indicate the loadings of each trait in the PCA. The legend shows the amount of functional space (i.e., functional richness) occupied at 50% and 99% probabilities (indicated by thick contour lines). The red regions falling within the limits of 50% probability correspond to the hotspots mentioned in the text. wea, weaning length; fmat, time to reach female maturity; ls, size of litter; gest, gestation length; long, longevity; ly, number of litters yearly. **b – d,** Patterns in space occupation of aerial (**b**), terrestrial (**c**), and aquatic (**d**) mammals. Legends show functional redundancy of the realm calculated as absolute values of the Standardized Effect Size (Red_99%_). SES was calculated considering the amount of functional space occupied by observed and randomized realm at 99% probability. * is SES p-value < 0.05; ** SES p-value < 0.01; *** SES p-value < 0.001. In all panels, the colour gradient (red, orange, and white) depicts different density of species in the space (red areas have higher density of species). Silhouettes of characteristic species from each realm were downloaded from PhyloPic (http://www.phylopic.org/).

## Methods

### Data collection and trait imputation

To characterise the mammals’ life history strategies, we used the Amniote database which comprises information for ca. 18,300 species of mammals, reptiles, and birds (Myhrvold *et al*. 2015). We extracted mammalian species (N = 4,953), and selected the life history traits with the most complete information *(i.e.,* traits with records for at least 1,000 species): litter size (ls, number of offspring per reproductive event), litter per year (ly, number of litters per year), age at female maturity (fmat, time needed for female individuals to reach maturity in days), weaning length (wea, time needed to wean the offspring in days), gestation length (gest, time passing between conception and birth in days), longevity (long, individual lifespan in years) and adult body mass (bm, in grams). For finer definition and information of traits’ measurements please refer to Myhrvold et al. (2015).

Because no trait was fully informed for all species, we imputed missing life history trait values. Recent studies have shown that imputation of individual traits allows to accurately characterize the position of species in trait spaces that are the result of combining different correlated traits (see Carmona, Tamme, et al. 2021; Carmona, Bueno, et al. 2021 for examples). Additionally, recent simulations indicate that examining patterns in space occupation built considering imputed species, produces results much closer to real patterns in space occupation compared to considering only species with complete trait information (Stewart *et al*. 2023). Because accounting for evolutionary relationships improves the accuracy of the imputation (Penone *et al*. 2014), we downloaded mammalian phylogenies from Vertlife database (Upham *et al*. 2019). We selected the complete fossil-based set of 10,000 phylogenies and computed a single maximum clade credibility tree (MCC tree) using the *maxCladeCred* function of *‘phangorn’* R package (Schliep 2011). The 135 species absent in the phylogeny but with life history information were added at the genus’ root using *add.species.to.genus* function of *‘phytools’* R package (Revell 2012). Species that were missing in the life history traits database were pruned using *drop.tip* function of *‘phyltools’* R package.

For the imputation procedure, we selected mammalian species with information for body size and at least one of the other considered life history traits (*i.e.,* ls, ly, fmat, wea, gest, long), resulting in a subset of 3,438 species. Within this subset 1,293 species had complete trait measurements (c.a. 37.6% of the total). Then, following Carmona, Tamme, et al. 2021, we included phylogenetic information in the imputation procedure using the first 10 phylogenetic eigenvectors derived from the MCC tree. For the imputation, we used the *missForest* function from the *‘missForest’* R package(Stekhoven 2022). After the imputation, we estimated the reliability of the imputation procedure (Supplementary methods 1). The imputation procedure efficiently retrieved the real position of a species, with an average error of 2.8% of the range for the first component and 1.7% for the second component. Imputations errors were consistent across orders with different level of data completeness and coverage.

The final procedure resulted in a dataset of 3,438 mammalian species (53.7% of currently recognized living mammals, Burgin *et al*. 2018) from 29 orders in which 37.6% of traits information (1,293 species) derived from empirical measurement and 62.4% (2,145 species) were imputed.

### Ascribing species to environmental realms and mode-of-life

In order to have information on species adscription to the terrestrial, aerial, and aquatic realms, we downloaded information on the environment exploited by each mammal from the Worms (WoRMS Editorial Board 2020) and IUCN (IUCN 2017) databases using the R packages *‘worms’*(Holstein 2018) and *‘redlist’* (Chamberlain 2020), respectively. We classified mammalian species as “terrestrial” (*i.e.,* land realm) in case of unambiguous classification from both databases, whereas we limited “aerial” category (air) to powered flight species (*i.e.,* Chiroptera order). Although there is no doubt about the classification of “fully” aquatic species (*i.e*., species that spent their entire life in water, *e.g.,* dolphins, whales, dugongs, Perrin *et al*. 2009), there is not a strict classification of semi-aquatic species (Hood 2020). Due to this, the Worms and IUCN databases classification did not match for many species. For instance, the rhesus monkey, *Macaca mulatta*, is classified as semi-aquatic according to the Worms database, while IUCN classifies it as terrestrial. To solve this problem, we performed a finer classification of aquatic species by summarizing species’ dependency on water. First, we selected all species classified as freshwater and marine in at least one of the IUCN and Worms databases. Then, we created an Aquatic Dependency Index (AD Index) to classify species as “fully” aquatic, “fully” terrestrial, or semi-aquatic. We built the AD Index answering a series of binary questions capable of summarizing the amount of time spent in water during fundamental phases of species life histories (*e.g.,* birth, mating; Supplementary methods 3). The AD Index was used to refine aquatic classification by relabelling species scoring zero in the AD Index (*i.e.,* no important parts of life are strictly constrained by water presence) as terrestrial species. Then, we classified as aquatic (hereafter aquatic) all species scoring 5 out of a maximum of 5 on the AD Index (*i.e.,* species that spent their whole life in water, like dolphins), while we classified as semi-aquatic all species scoring between 1 and 4 (Supplementary methods 3). Out of the 3,438 mammalian species considered in our analyses, 2,634 species were classified as terrestrial (76.6% of total set of species), 661 species were classified as aerial (19.2% of the total), 85 species were classified as aquatic (2.5% of the total), and 58 species were classified as semi-aquatic (*i.e.,* transitional species, 1.7% of the total).

We further classified terrestrial species according to their arboreal, semiarboreal, and fossorial mode-of-life by using the same classification applied in Santini, *et al*. 2022. We manually added missing information throughout a literature review(Bels & Russell 2023; IUCN 2017; University of Michigan 2022). Out of the 2,634 species classified as terrestrial, 595 species were classified as arboreal (22.6% of terrestrial species), 175 species were classified as semi-arboreal (6.6% of terrestrial species), 117 species were classified as fossorial (4.4% of terrestrial species), and 1,747 species were classified as ‘fully’ terrestrial (*i.e.,* confined on land, 66.3% of terrestrial species).

Species taxonomies across all previously mentioned data tables (*i.e.,* life history traits, phylogeny, Worms, IUCN, and mode-of-life databases) were standardized using *gnr_resolve* function of the *‘taxize’* R package (Chamberlain & Szöcs 2013) resolving for the Global Biodiversity Information Facility (GBIF) Backbone Taxonomy.

### Construction of the global life history traits space

Body mass is a strong allometric constraint that affects the values of many other life history traits (Bielby *et al*. 2007; Jeschke & Kokko 2009; Sibly & Brown 2007). Mammals’ body mass can span from a few grams of shrews to tons for whales, and the aquatic realm strongly selects for bigger sizes compared to the terrestrial realm (Gearty *et al*. 2018). Therefore, to address life history strategies without confounding effects from different orders of magnitude, we accounted for the effect of body mass before computing the mammalian life history space. For this, we performed a series of ordinary linear regressions between single log-transformed life history traits and log-transformed species’ adult body mass. Then, we extracted body mass–corrected residuals for each species from each regression (Jeschke & Kokko 2009; Revell 2009). Body mass–corrected residuals were further centred, scaled, and used as size-corrected life history traits (Supplementary material Fig. 1).

Mapping species’ position in multidimensional spaces based on trait information allows the identification of major axes of trait variation (Bueno *et al*. 2023; Díaz *et al*. 2016), as well as summarizing species’ ecological strategies. We constructed the mammalian life history space by performing a principal component analysis (PCA) on the complete set of mammalian species. We retained the first two axes which together account for 75.1% of variance in mammalian life history traits.

Then we estimated the reliability of our methodological choices by comparing this life history space (*i.e.,* the scores of species in the selected number of principal components) with three other spaces: one built considering only complete set of records (*i.e.,* non-imputed data; N = 1,293; Supplementary material Fig. 2); one built considering the imputed dataset without body mass correction (N = 3,438); and one built considering the same body mass–corrected residuals used to build the life history space (N = 3,438) but accounting for the phylogenetic history of species (Supplementary material Fig. 3; Supplementary material Table 1). In all abovementioned cases, the comparisons revealed strong correspondence between spaces, demonstrating that our methodological choices do not influence the inferred relationships between life history traits (Supplementary Methods 2).

### Exploring patterns within the global life history space

We derived life history structure (*i.e.,* the patterns of organization of species in the life history space) using the trait probability density (TPD) framework (Carmona *et al*. 2016), which computes species’ probabilistic distribution in the life history space using multivariate kernel density estimation. The kernel density estimation for each species was estimated as a multivariate normal distribution cantered at the species’ coordinates in the PCA space (Carmona *et al*. 2016, 2019), and with a standard deviation computed using the unconstrained bandwidth selectors from the *‘Hpi’* function in the *‘ks’* package (Duong 2007, 2022). To estimate the life history structure of the whole set of mammals, all species’ kernels were weighted and aggregated to form a single continuous TPD function across the whole mammalian life history space. The value of the TPD function at any given point of the life history space reflects the density of species in that particular area of the life history space (Carmona *et al*. 2016, 2019). We graphically represented the so-built mammalian life history structure by highlighting the contours containing 50% and 99% of the total probabilistic distribution of species (Fig. 1a). We used the *‘TPD’* R package (Carmona 2019; Carmona *et al*. 2019).

We used mammalian global structure to test whether mammals distribute in the life history space following a random assembly of life history traits (*i.e.,* life history strategies are randomly selected) or whether certain life history strategies are more prevalent than others (Carmona *et al*. 2021a; Díaz *et al*. 2016; Southwood 1988). First, we computed functional richness (the amount of functional space occupied (Carmona *et al*. 2016; Mason *et al*. 2005) at increasing probability thresholds (*i.e.,* from 0.1% to 99.9% of the TPD function, using 0.1% increments). In this way, we estimated a “profile” of the probabilistic distribution of species, reflecting what amount of the life history space is occupied at different probability thresholds. We compared the observed functional richness profile with a null model (999 repetitions) considering a bivariate normal distribution of species with the same mean and variance as the observed data (see Carmona, Bueno et al., 2021). This analysis was performed by sampling 3,438 simulated species *(i.e.,* complete set of mammals) from a bivariate normal distribution with the same mean and covariance of observed data calculated using the ‘*rmvnorm’* function of the R package *‘tmvtnorm’*(Wilhelm & Manjunath 2010). In the same way as done for the observed data, for each repetition of the null model we computed life history structure and measured functional richness at increasing probability thresholds. We then compared the observed and null functional richness profiles; functional richness values lower than expected under the null model would suggest that the global set of mammals occupy a smaller portion of the life history space than expected under a normal distribution and vice-versa.

Then, using the same null model, we tested whether the portions of life history space that are mainly exploited by mammalian species are closer or farther than expected from the centre of the distribution. For this, we estimated functional divergence (*i.e.*, degree to which species’ density is skewed towards the extremes of the life history space) considering 99% of the total species’ probability (Carmona *et al*. 2016; Mason *et al*. 2005) for both observed and simulated data. High functional divergence would reveal that species are segregated towards the edges of the life history space, whereas low divergence would indicate that species are clustered in the centre of the life history space (Carmona *et al*. 2021a).

We estimated the standardized effect size (SES) of both the functional profiles (at each probability threshold) and the functional divergence measures as: SES = (observed value – mean (simulated values))/standard deviation (simulated values). SES values express the number of SD units by which the observed value deviates from the mean of the simulated ones. A SES value lower than 0 indicates that the observed metric is smaller than the average of the simulated values and vice versa. Finally, we tested the statistical significance of observed and simulated differences by estimating two-sided p-values computed by confronting the SES values with a cumulative normal distribution with mean = 0 and standard deviation = 1 (Carmona *et al*. 2021a, Supplementary material Fig.4).

Lastly, to have a clearer picture of how different species are positioned in the life history space, we computed TPD functions for each mammalian order with at least 6 species (total of 18 orders). We graphically represented the mammalian life history structure of each order by highlighting the contours containing 50% and 99% of the total probabilistic distribution of species order (Supplementary material Fig. 5).

### Exploring environmental realms’ patterns in the life history space

We produced realm-specific TPD functions by aggregating the species that are characteristic of each realm. We followed the same procedure explained above for the global set of mammalian species. In the same way as done with the complete mammalian life history structure, we represented graphically the TPD for each realm and highlighted contours containing 50% and 99% of the total probability (Fig. 1b-d).

We compared the structure of each of the environmental realms (*i.e.,* air, land, water and transitional) considering multiple aspects. First, we compared realms in terms of the amount of life history space that they occupy (*i.e.,* their functional richness). Because functional richness is known to be positively related to the number of species present in a group, we built a null model (999 repetitions) randomizing species assignment (Carmona *et al*. 2021a). For each realm, we randomly extracted the same number of observed species from the complete mammalian database and computed TPD functions of the random community. We computed functional richness at the 99% of total probability for both observed and randomized structures and calculated SES as explained above. Under this null model, negative SES values would indicate that the species present in the realm are more clustered in the life history space (*i.e.,* occupy a smaller portion of the life history space, and therefore species are redundant in their life history strategy) than expected given the same number of randomly selected species and vice versa. A highly clustered realm would indicate high redundancy of life history strategies (Carmona *et al*. 2021a), suggesting that the prevalent conditions in that realm exert a strong selection on life history strategies. This procedure allows to compare functional richness among realms with very different number of species (Supplementary material Table 2a).

Then, we statistically tested whether dissimilarities in species positioning were a product of a realm-driven selection of life history strategies. We estimated how much of the total variation in the position of the species on life history space was explained by difference in realms. For this, we first calculated the dissimilarities (using Euclidean distances) between all pairs of species in the life history space. Then, we performed a PERMANOVA analysis considering the three major environmental realms (*i.e.,* air, land, and water) as the explanatory variable (R package *‘vegan‘* (Oksanen *et al*. 2020)). If the realm explains a large portion of life history variation, it means that life history variability is mainly related to differences between realms rather than differences among species within the realm (Carmona *et al*. 2021a).

We estimated pairwise dissimilarities in life history structures between pairs of realms (*i.e.,* air, land, water, and transitional, Fig. 2). Dissimilarities were calculated as 1 - overlap of the corresponding TPD functions producing measures varying between 1, indicating a complete overlap between the considered TPD functions, and 0, when the two functions are completely disjoint. TPD-based dissimilarities consider both differences in the boundaries and differences in the density of the occupation of the life history space (Carmona *et al*. 2021b; Germain *et al*. 2023; Toussaint *et al*. 2021). These dissimilarities can be further decomposed into two complementary components: one quantifying up to what point the life history differences between realms are related to the occupation of exclusive regions of the space (*i.e.,* turnover) and another reflecting how differently the two realms occupy the shared parts of the life history space (*i.e.,* nestedness Carmona *et al*. 2019).

**Fig. 2.**
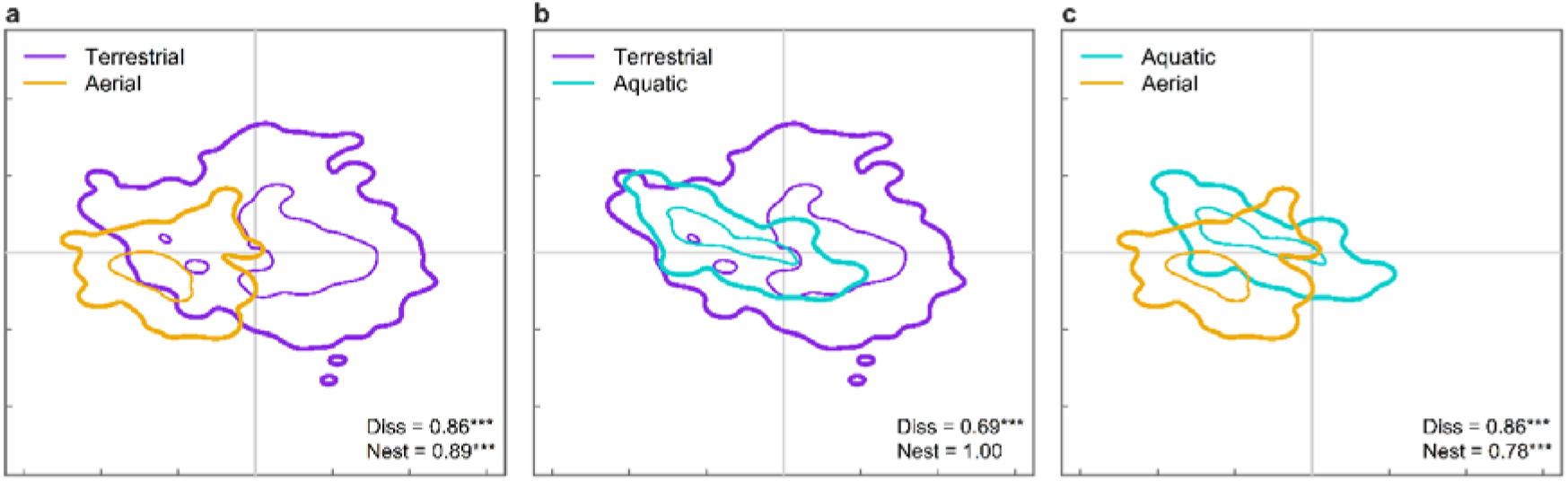
Environmental realms select for distinct yet nested life history strategies. Overlap-based dissimilarities between terrestrial (purple), aerial (light orange), and aquatic (light blue) species distributions in the life history space. Specifically: **a,** dissimilarity between terrestrial and aerial realms; **b,** dissimilarity between terrestrial and aquatic realms; **c,** dissimilarity between aerial and aquatic realms. Each realm is highlighted at 50% (fine lines) and 99% (thick lines) of total probability. SES of dissimilarity and nestedness values for each pairwise comparisons are reported in the legend together with SES p-values (* SES p-value < 0.05; ** SES p-value < 0.01; *** SES p-value < 0.001).

We tested whether the observed dissimilarities between environmental realms were significantly higher than expected by chance by means of null models (Micó *et al*. 2020; Traba *et al*. 2015). In this null model, for each pair of realms, we considered only the set of species belonging to both realms and we randomized the realm classification. We maintained the same number of species observed in each considered realm and avoided species assignment to multiple realms. For each repetition of the null model (n = 999), we estimated the TPD functions of each realm and estimated their dissimilarity, as well as its nestedness and turnover components. Finally, we computed SES values for all mentioned metrics as described above (Supplementary material Table 2b).

Lastly, we explored whether dissimilarities between environmental realms were maintained also in closely related species. To do this, we computed TPD functions for orders having species inhabiting in different realms. Because all aerial species belong to the Chiroptera order, we decided to focus on orders having both terrestrial and aquatic species. We extracted orders that had at least six species in both the aquatic (in this case identified by both aquatic and semi-aquatic species) and terrestrial realms. For each of these orders, we calculated a life history structure for terrestrial species and one for aquatic species. We represented graphically structures highlighting 50 and 99% of the total probability and we computed pairwise dissimilarities between the terrestrial and aquatic life history structure of each order (Supplementary material Fig. 6).

### Extracting differences in terrestrial mode-of-life and species life histories

For the terrestrial realm, we further produced specific TPD functions for each of the considered mode-of-life (*i.e.*, arboreal, semiarboreal, fossorial, and confined on land) following the same procedure as above. We represented graphically the TPD for each mode-of-life and highlighted contours containing 50% and 99% of the total probability (Supplementary material Fig. 7). As above, we computed observed functional richness for each mode-of-life, and we performed a null model to extract SES values of functional richness (Supplementary material Table 3a). Then, we estimated pairwise dissimilarities in life history structures across all different mode-of-life and environmental realms (*i.e.*, aerial, aquatic, transitional) following the same procedure as above (Supplementary material Fig. 8). We tested whether the observed dissimilarities were significantly higher than expected by chances by performing the same null model as done above for environmental realms (Supplementary material Table 3b).

We tested whether, within taxonomically related species, exploiting the environment with different mode-of-life allows for distinct set of life history strategy than species confined on land. For that, we selected all mammalian orders showing arboreal, semiarboreal, or fossorial species and species confined on land. Within each order, we extracted the centroid of the functional structure of the species confined on land as well as the centroid for the functional structure of the mode-of-life exploited in the order (Fig. 4). Then, we produced three datasets (one for arboreality, one for semi-arboreality, and one for fossoriality), containing the centroids the behaviours together with the ones of species confined on land for each of the considered order. We used these datasets to statistically test for difference between species confined on land and species showing different mode-of-life through a series of linear mixed effect models using the R package *‘lmerTest ‘* (Kuznetsova *et al*. 2017). For each dataset we performed two models, one for each principal component, using the mean score of the order along the PCs as response variables, mode-of-life as fixed effect, and mammalian orders as random intercept (Supplementary material Table 4).

Similarly to arboreal, semiarboreal, and fossorial mode-of-life, species with high encephalisation are capable to exploit the environment in a plastic way, avoiding hazards and stressful situations (González-Lagos *et al*. 2010; Zhu *et al*. 2023). Consequently, we further explored whether high encephalization (Barton & Capellini 2011; González-Lagos *et al*. 2010; Pérez-Barbería *et al*. 2007; Reader & Laland 2002) allows to exploit portion of the life history space otherwise not successful for the terrestrial realm (Supplementary methods 4). The final procedure resulted in two generalized additive models (GAMs) mapping species observed brain mass (Barton & Capellini 2011; Benson-Amram *et al*. 2016; brain mass information were extracted from Combine database Soria *et al*. 2021, for a subset of 1,054 species with non-imputed brain mass observations) in response to the species position in the life history space (Supplementary material Fig. 9).

All statistical analyses were performed using R version 4.0.3 (R Core Team 2020).

### Results & Discussion

### Global life history strategies are constrained and profoundly divergent

Mammalian life history traits vary mostly along a two-dimensional space, which suggests a strong trait covariation limiting viable life history strategies. The first two principal components (PC) of the PCA account for 75.1% of life history variation, conforming a life history space characterised by two primary trade-offs: litter size *vs.* gestation length, and number of litters per year *vs.* age at female maturity and weaning length (Fig. 1a). The configuration of this space is line with previous studies on smaller sets of species (267 species in Bielby et al., 2007; 80 species in Gaillard et al., 1989). The first dimension of this space (PC1) captures the fast-slow continuum (Stearns 1992), distinguishing fast species with short lives, gestation and weaning, early maturity, and frequent litters, from slow species with fewer and infrequent litters, longer lives, and later maturity (Fig. 1a). The second dimension (PC2) reflects lifetime reproductive effort, illustrating the trade-off between litter size and frequency, as well as offspring quality (*i.e.,* altricial *vs.* precocial), and reproductive frequency (*i.e.,* semelparity *vs.* iteroparity). Species’ positioning and life history trait correlations in the life history space remain consistent after accounting for evolutionary history (Procrustes test r = 0.95; p < 0.001 for species; angles correlation r = 0.85, p < 0.001 for traits; see Supplementary methods 2), suggesting that this two-dimensional life history strategy space applies to all mammals, independent of shared ancestry (Supplementary material Fig. 3).

Mammalian species exhibit a remarkably uneven, concentrated distribution within the life history space. Comparing global mammal occupation patterns along this space to null models representing bivariate normal distributions with identical mean and covariance (Carmona *et al*. 2021a), the occupied space is smaller than expected by chance (Supplementary material Fig. 4a). For example, at the 99% quantile of species probability distribution, observed functional richness (*i.e*., amount of space occupied by species) is 32% smaller than the simulated counterpart (SES_Fric99%_ = -11.79, p < 0.001, n = 999 iterations). Additionally, species’ distribution is much more divergent than the simulated (SES_Fdiv_ = 18.79, p < 0.001; n = 999 iterations; Extended data Fig.4b), primarily due to species clustering in two hotspots at opposite ends of the fast-slow continuum. Our findings reveal a dichotomy in successful investments to survival, development, and reproduction of global mammals: species investing on quantity of reproductive investment contrast with those prioritizing quality (Grime & Pierce 2012; Southwood 1988; Stearns 1992). Species in the “fast hotspot” (Fig. 1), such as mice and rabbits, optimize fast turnover and high reproductive output, enabling them to propagate their genes in unpredictable and unfavourable environments (Grime & Pierce 2012; Southwood 1988). Conversely, “slow hotspot” species (Fig. 1a), including primates, cetaceans and bats, typically make costly parental investments to equip their few offspring with advantageous adaptations for long-term persistence and environmental exploitation (Barton & Capellini 2011; Grime & Pierce 2012; Southwood 1988). This bimodal pattern mirrors that of plant traits (Carmona *et al*. 2021a; Salguero-Gómez *et al*. 2016), suggesting that, across the tree of life, species tend to cluster around a few contrasting syndromes that lead to successful strategies (Junker *et al*. 2022).

### Divergent life history strategies correspond to different environmental realms

Major environmental realms select for distinct and highly redundant life history strategies (Fig. 1b-d; Fig. 2). This finding is confirmed by a PERMANOVA analysis, which attributes 35% of variation in life history traits to differences among realms. The marked differences in life history strategies among realms are further demonstrated by the dissimilarities between pairs of realms (D_land-water_ = 69%, D_land-air_ = 86%, D_air-water_ = 86%; p < 0.001 in all cases; Fig. 2; Supplementary material Table 2), as well as by the high levels of species redundancy within individual realms (*i.e.,* species within an environmental realm tend to have similar trait values; R_land_ = 7.99, R_air_ = 9.25, R_water_ = 4.38; p < 0.001 in all cases; Supplementary material Table 2). Terrestrial species disproportionally occupy the “fast hotspot” (Fig. 1c), including representatives of marsupials as well as the Carnivora, Lagomorpha, Rodentia, and Eulipotyphla orders (Supplementary material Fig. 5). Primates and Perissodactyla are the only exceptions to this pattern since they cluster in the slowest portion of the life history space. On the contrary, aerial (Fig. 1a) and aquatic species (Fig. 1d) are confined to the slow end of the spectrum. Despite the similar pace of life of aquatic and aerial species, their reproductive strategies differ substantially. Aerial species exhibit long gestation times, while aquatic species display late female maturity and extended weaning (Fig. 1a). The predominance of slow strategies in aquatic species contrasts with previous observations mainly based on baleen whales (Gaillard *et al*. 1989), which are among the longest-lived aquatic species (Fig. 1d). Our study, however, includes dolphins, displaying a more complete picture. Dolphins and baleen whales exhibit distinct migratory, dietary, and social habits (Perrin *et al*. 2009), influencing resource perception and gathering within realms and affecting life history variation (Famoso *et al*. 2018; Soriano-Redondo *et al*. 2020; Zhu *et al*. 2023).

Despite marked differences and redundancy among realms, characteristic strategies of a given environmental realm can be observed in others. This shared set of life history strategies is particularly evident in the case of the aquatic realm, fully contained within the life history space occupied by terrestrial species (Nestedness_land-water_ = 1, Fig. 2b). The aerial realm is also highly overlapping with the terrestrial one (Nestedness_land-air_ = 0.89; Fig. 2a); primates, a small subset of terrestrial species, are the main reason for such an overlap. For example, apes (including humans), and dolphins have essentially the same life history traits, whereas terrestrial Lemuriformes and Tarsius species are located in the part of the life history space occupied by bats (Figure 1; Supplementary material Fig. 5).

### Species transitioning between realms have intermediate life history strategies

The life history strategies of semi-aquatic species provide further evidence for the profound influence of environment realms on life history variation. Semi-aquatic species exhibit intermediate life history strategies, slower than terrestrial species but faster than fully aquatic ones (Fig. 3). This pattern holds within semi-aquatic mammals, whereby species occupy different regions of the life history space depending on their degree of water-dependence. For example, species with greater aquatic involvement, such as seals and walruses (Davis 2019; Perrin *et al*. 2009), occupy portions of the life history space similar to fully aquatic species like manatees, beluga, or sperm whales. Conversely, species with lesser water-dependence, like rodents and otters (Davis 2019; Hood 2020; Perrin *et al*. 2009), are located in the hotspot typical of terrestrial mammals (Fig. 3). Interestingly, redundancy of semi-aquatic mammals was not significantly different from what expected by chance (R_semi-aquatic_ = -0.89; p = 0.27; Supplementary material Table 2). We advocate for future research to investigate the evolution of aquatic and semi-aquatic adaptations, in order to understand the realized diversity of the mammalian return to water (Davis 2019).

**Fig. 3.**
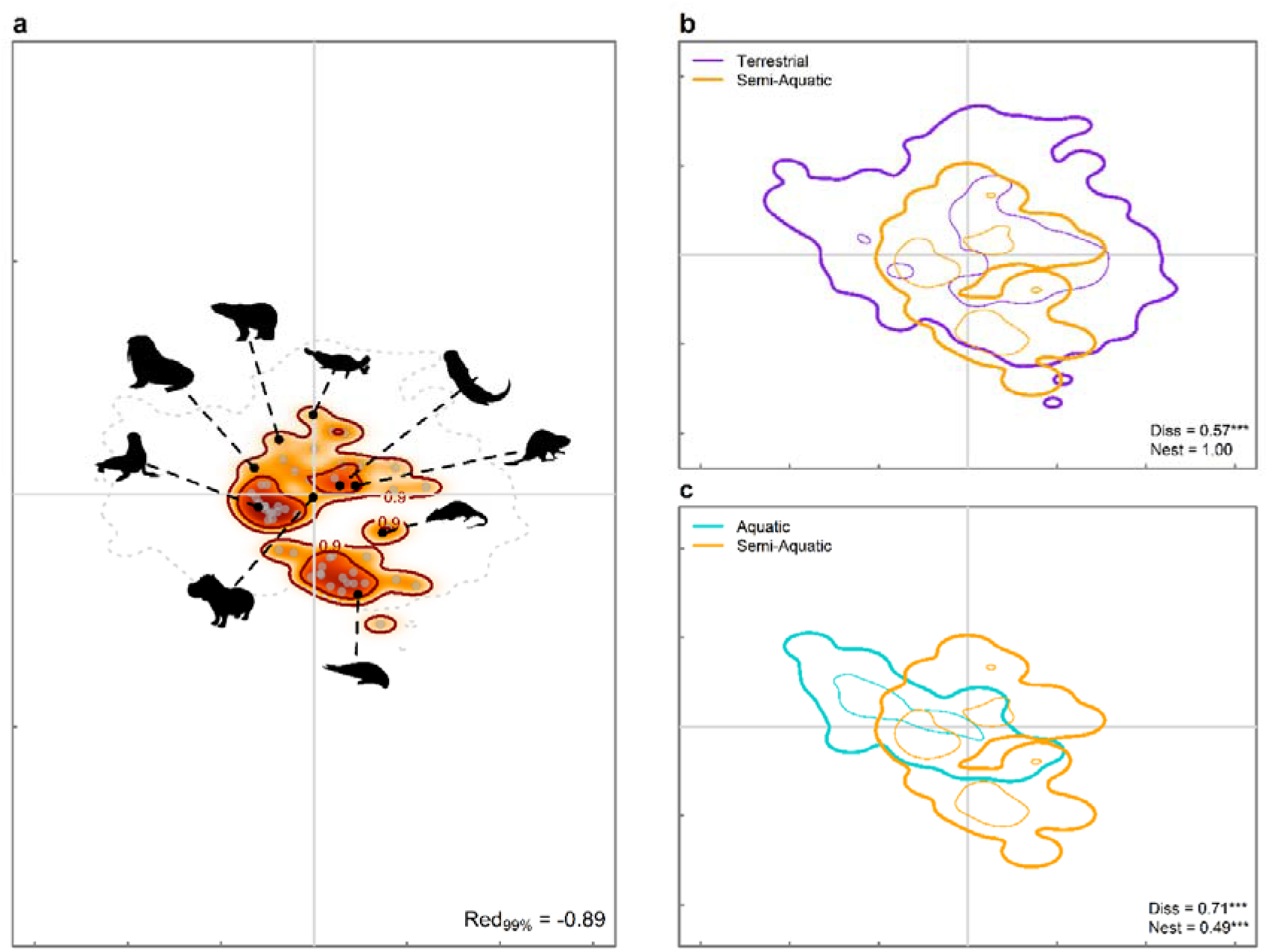
Semi-aquatic species transition between aquatic and terrestrial life histories depending on the evolutionary time they spent in water. **A,** Occupation of transitional mammals in the life history space. The colour gradient (red, orange, and white) depicts different density of species in the space (red areas have higher density of species). The legend shows the amount of functional space (*i.e*., functional richness) occupied at 50% and 99% probabilities (indicated by thick contour lines). Dashed grey line show the portion of space occupied by global mammals. Legend show functional redundancy of transitional species calculated as absolute values of the Standardized Effect Size (Red_99%_). SES was calculated considering the amount of functional space occupied by observed and randomized realm at 99% probability. Silhouettes of characteristic semi-aquatic species were downloaded from PhyloPic (http://www.phylopic.org/). **B-c,** pairwise overlap-based dissimilarities between semi-aquatic (SA) and both terrestrial (**b**) and fully aquatic (**c**) mammals. In both panels, contours at 50% (fine lines) and 99% (thick lines) are highlighted for terrestrial (purple), aquatic (light blue), and semi-aquatic (orange) species. Legends show observed dissimilarities between realms and semi-aquatic species (Diss) and value of nestedness (Nest). P-values for the standardized effect size (SES) of dissimilarity and nestedness are present only for significant SES. ***: SES p-value < 0.001. All panels show the probabilistic species distributions in the spaces defined by the two first principal components (PC1 and PC2) of PCA considering life history traits for all mammals.

The reported intermediate life history strategy location of semi-aquatic mammals is independent of taxonomic order. This robustness confirms the association between environmental realms and life history strategies even among closely related species. For example, terrestrial and semi-aquatic carnivores separate in the life history space following the same patterns seen in the global dataset with terrestrial carnivores being faster than their semi-aquatic counterparts (Supplementary material Fig. 6a-c). Moreover, the sea otter (*Enhydra lutris*), the only fully aquatic carnivore, exhibits a markedly different life history strategy compared to semi-aquatic otters (Supplementary material Fig. 6). Among Cetartiodactyla, aquatic (dolphins, whales) and semi-aquatic (hippos) species have slower life history strategies than terrestrial species (giraffe, bison, gazelles; Supplementary material Fig. 6d-f). Altogether, these findings suggest that species transitioning between aquatic and terrestrial realms possess intermediate life history strategies.

### Slow life history strategies in arboreal and fossorial species

Arboreal, semiarboreal, and fossorial mode-of-life are associated with distinct life history strategies than land-confided species. Species confined on land mainly clustered in the fast portion of the life history space, coinciding with the “fast hotspot” of mammalian global structure (Supplementary material Fig. 7c). Conversely, arboreal, semiarboreal, and fossorial species mainly cluster towards the slow (arboreal, Supplementary material Fig. 7a) and intermediate (fossorial and semiarboreal; Supplementary material Fig. 7b-c) portions of the life history space. These discrepancies between land-confined species and those using specific mode-of-life to dwell in the terrestrial realm are confirmed by dissimilarities across pairwise comparisons (D_land-arboreal_ = 61%, D_land-semiarboreal_ = 47%, D_land-fossorial_ = 53%; p < 0.001 in all cases) and hold consistent across taxonomically related species (Fig. 4). Arboreal and fossorial species are significantly skewed towards the slowest portion of the life history space in comparison to representatives of the same order confined on land. Conversely, the pattern for semiarboreal species is less clear and fluctuates among orders (Fig. 4, Supplementary material Table 4), suggesting a role of arboreal (and fossorial) specialization in the selection of slow life history strategies on land.

**Fig. 4.**
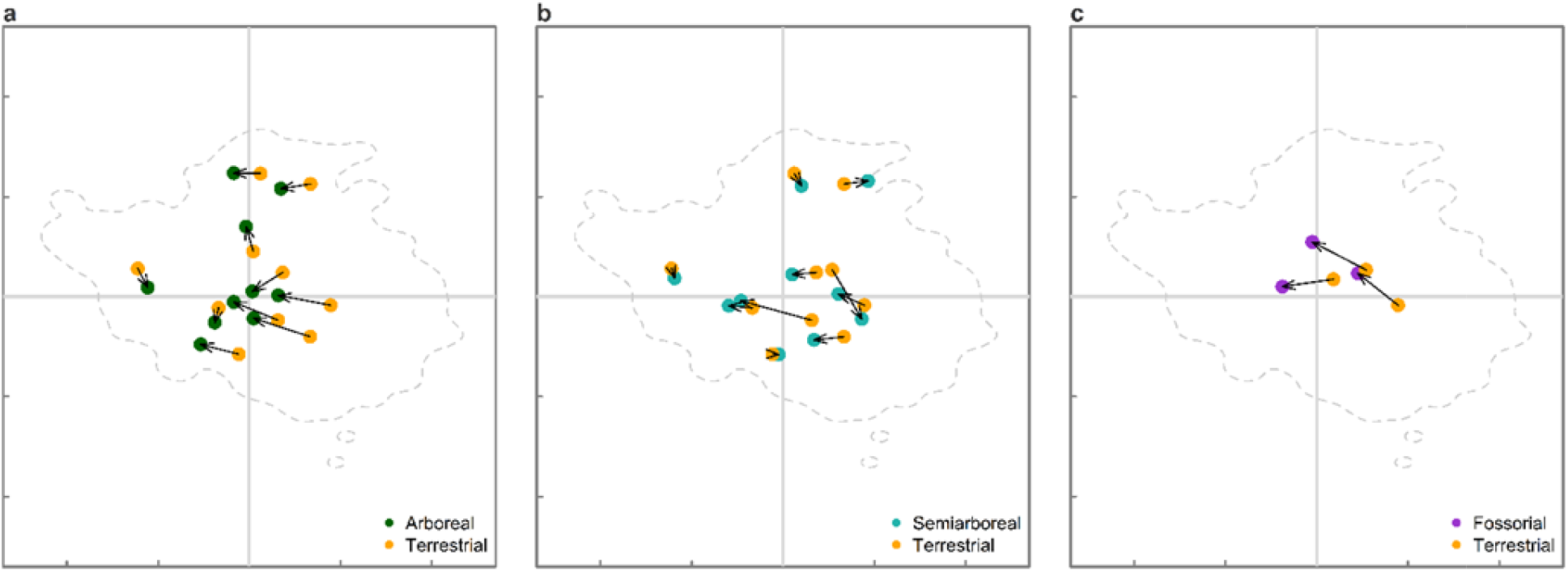
Arboreal and fossorial mode-of-life are associated with slower life histories in taxonomically related terrestrial species. Differences in centroid among the life history structures of species confined on land (terrestrial) and **(a)** arboreal, **(b)** semiarboreal, and **(c)** fossorial species of the same order. Each arrow connects the centroid of the terrestrial structure and mode-of-life structure present within the same order. Green dots are arboreal centroids, aquamarine dots are semiarboreal centroids, purple dots are fossorial centroids, and orange dots are centroids of species confined on land.

Arboreal and semiarboreal adaptations expand the portion of the life history space occupied by the terrestrial realm towards life history strategies that are otherwise exclusive of the aerial and aquatic realms. The overlap we detected between terrestrial realm and both aerial and aquatic realms (Fig. 2) is primarily related to the overlap with arboreal and, to a lesser extent, semiarboreal life history strategies (Supplementary material Fig. 8). Indeed, terrestrial and aerial nestedness decreased by 55% (increasing the dissimilarity between the two realms by 10%) when considering only terrestrial species confined on land despite these species being 66.3% of terrestrial species. The dissimilarity between aquatic and land-confined species increases by 15% if compared to the dissimilarity computed considering the full set of terrestrial adaptation (Supplementary material Table 3b). Although with different extent, arboreal, semiarboreal, and fossorial mode-of-life, allow species to escape mortality risks as well as to explore different ecological opportunities (Bels & Russell 2023; Scheffers *et al*. 2017; Stroud & Losos 2016; Withers *et al*. 2016). Similarly to the aquatic and aerial realms, the access of vertical movement may allow species to modify how the environmental pressures are perceived in the terrestrial realms allowing for different life history strategies otherwise unfavoured on land (Greenslade 1983; Grime & Pierce 2012).

Arboreal species exploit life history strategies that are characteristic of the aquatic realm (Supplementary material Fig. 8). Interestingly, dissimilarities between arboreal and aquatic species are lower than the one observed between arboreal and land-constrained species (D_water-arboreal_ = 55%, D_land-arboreal_ = 61%, p < 0.001). Accordingly, the most prevalent aquatic life history strategies (50% of the probability distribution) are completely nested within arboreal ones, overlapping in the portion of the life history space occupied by arboreal primates and dolphins (Fig. 1,2a; Supplementary material Fig. 5,8). This shared set of life history strategies among primates and dolphins, might be linked with complex social structures and high encephalization in both groups (Perrin *et al*. 2009; Ridgway *et al*. 2017; Worthy & Hickie 1986). Similarly to arboreal, semiarboreal, and fossorial mode-of-life, high encephalization, which generally relates to both social and behavioural facets (Barton & Capellini 2011; Pérez-Barbería *et al*. 2007), allow species to exploit the environment in a plastic way, avoiding hazards and stressful situations (González-Lagos *et al*. 2010; Zhu *et al*. 2023). We tested whether high encephalization may play a role in selecting terrestrial strategies, by modelling the brain-mass ratio of terrestrial mammalian species (i.e. the ratio between brain dimension and body mass (Benson-Amram *et al*. 2016; Worthy & Hickie 1986) as a function of species’ position in the life history space (Supplementary methods 4). We found a robust association between larger brain mass and slow life histories (R^2^ = 0.48, Supplementary material Fig. 9), suggesting that larger brain size may significantly influence the diversity of terrestrial life history strategies. Consequently, while the environmental realm in which species live seems to drive life history strategies, behavioural characteristics of species seem to play a role in expanding the set of life history strategies exploitable in the terrestrial realm (Cox *et al*. 2021). Specifically, we suggest that vertical movement (*e.g.* arboreality, fossoriality) as well as plastic or social behaviours may allow species to modify the environmental pressures perceived under specific environmental constraints (Greenslade 1983; Grime & Pierce 2012; Southwood 1988), playing a fundamental role in defining the set of life history strategies realized under similar environmental requirements.

## Conclusion

Our analyses of over 3,400 mammal species reveal that major environmental realm is a strong predictor of mammalian life history strategies. Aerial (Babich Morrow *et al*. 2021; Gaillard *et al*. 1989) and aquatic (Davis 2019; Gearty *et al*. 2018; Uhen 2007) realms require specific adaptations, such as flight-capable skeletal structures (Barclay 1994; Maina 2000) or enhanced thermoregulatory abilities for aquatic life (Davis 2019; Perrin *et al*. 2009). These adaptations slow down and prolong lifespans in ways seemingly unexploited in the terrestrial realm by mammals (Babich Morrow *et al*. 2021). However, despite each realm’s unique set of life history strategies, all of them overlapped to some degree within the life history space due to species’ adaptations that enable the exploitation of otherwise unsuccessful strategies in each realm. Empirical research has shown that adaptations in the way species exploit the environment (Babich Morrow *et al*. 2021; Healy *et al*. 2014; Shattuck & Williams 2010), as well as social (Zhu *et al*. 2023), migratory (Soriano-Redondo *et al*. 2020), dietary characteristics (Famoso *et al*. 2018), and cerebral development (Barton & Capellini 2011; Seyfarth & Cheney 2002) can influence species’ life histories. Consequently, variations in these behavioural and ecological characteristics modulate species’ responses to environmental pressures (Grime & Pierce 2012) and expand the spectrum of life history strategies available for exploitation within a realm, as observed for activity patterns (Cox *et al*. 2021). For instance, differences in migratory, dietary, and social habits among baleen whales and dolphins (Perrin *et al*. 2009) may facilitate distinct life history strategies in aquatic environments, accounting for the rapid pace of life we observed in baleen whales. Here we demonstrate that arboreal, semi-arboreal and fossorial mode-of-life play a fundamental role in expanding the set of life history strategies in the terrestrial realms. Confirming empirical evidence (Babich Morrow *et al*. 2021; Bels & Russell 2023; Healy *et al*. 2014; Shattuck & Williams 2010), arboreal and fossorial mode-of-life favour slower life history strategies within taxonomically related species, with arboreal mode-of-life allowing terrestrial species to share life history strategies dominated by aquatic species. Similarly, the strong relationship we uncovered between brain dimension and the position of terrestrial species within the life history space (Supplementary material Fig. 9) suggest a role of sociality and encephalization in allowing terrestrial mammals to exploit the slower end of the life history spectrum. Accordingly, we advocate for future research to disentangle the influence of different ecological and behavioural adaptations on life history variation among terrestrial and aquatic mammals.

Our study provides a comprehensive assessment of global mammalian life history strategies, revealing the fundamental role of environmental realm in shaping these patterns. Remarkably, these patterns persist even among closely related species, highlighting the profound relationship between environmental realms and life history evolution. Our findings not only provide empirical insights into mammalian adaptation, but also establish a foundation for further exploration of ecological and evolutionary processes (Burgin *et al*. 2018; Grossnickle *et al*. 2019). In addition, this foundation may allow to improves our understanding of the drivers and consequences of ongoing global changes (Carmona *et al*. 2021b; Enquist *et al*. 2020) and future biodiversity patterns (Carmona *et al*. 2021b; Cooke *et al*. 2019; Toussaint *et al*. 2021). As we confront these challenges, our study emphasises the importance of understanding the intricate interplay between environment, ecology, and evolution to disentangle, anticipate, and mitigate the impacts on mammalian life history diversity.

## Acknowledgements

C.P.C and EB were supported by the Estonian Research Council grant (PSG293) and the European Regional Development Fund via the Mobilitas Pluss programme (MOBERC40). P.C.L. was supported by the European Union-Next Generation EU Maria Zambrano Program (ZAMBRANO 21-26). R. S-G. was supported by a NERC Independent Research fellowship (NE/M018458/1).

