## Extended data figure for "Global Diversity in Mammalian Life Histories: Environmental Realms and Evolutionary Adaptations"

**Supplementary material**

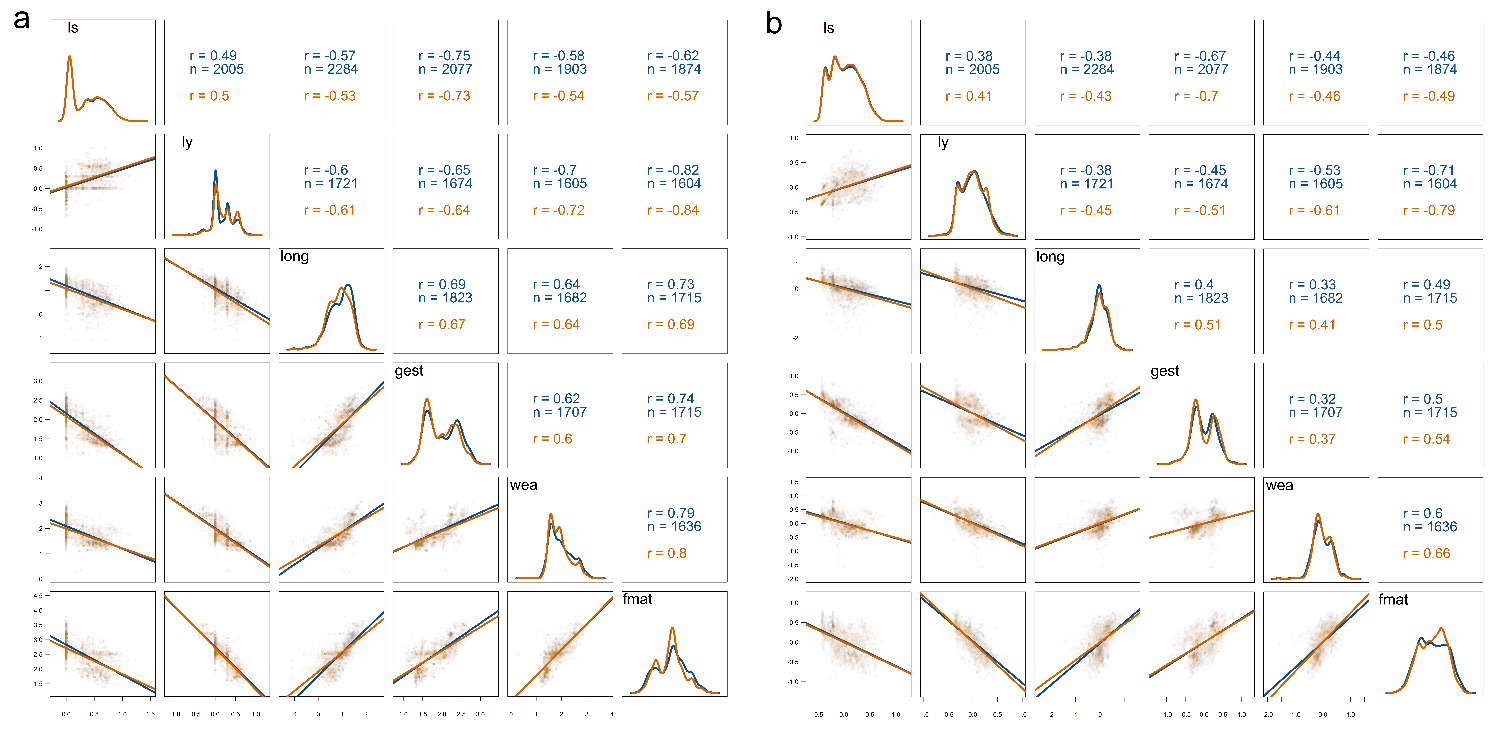

**Supplementary material Fig. 1 | Pairwise correlations between complete and imputed data. a,** complete and imputed observed traits; **b,** body mass residuals calculated from complete and imputed observed data. Both panels show pairwise correlations between life history traits in the different datasets (blue: complete dataset with 1,293 species, orange: imputed dataset with 3,438 species). The lower-left triangle of the matrix contains scatterplots of traits (after log-10 transformation) showing the relationship (including regression lines) between each pair of traits. The diagonal includes a probability density function showing the distribution of each individual trait. The upper-right triangle includes the value of the correlation coefficients and, in the case of the complete dataset, the number of species with empirical data for both traits (imputed dataset always considered the same numbers of species).

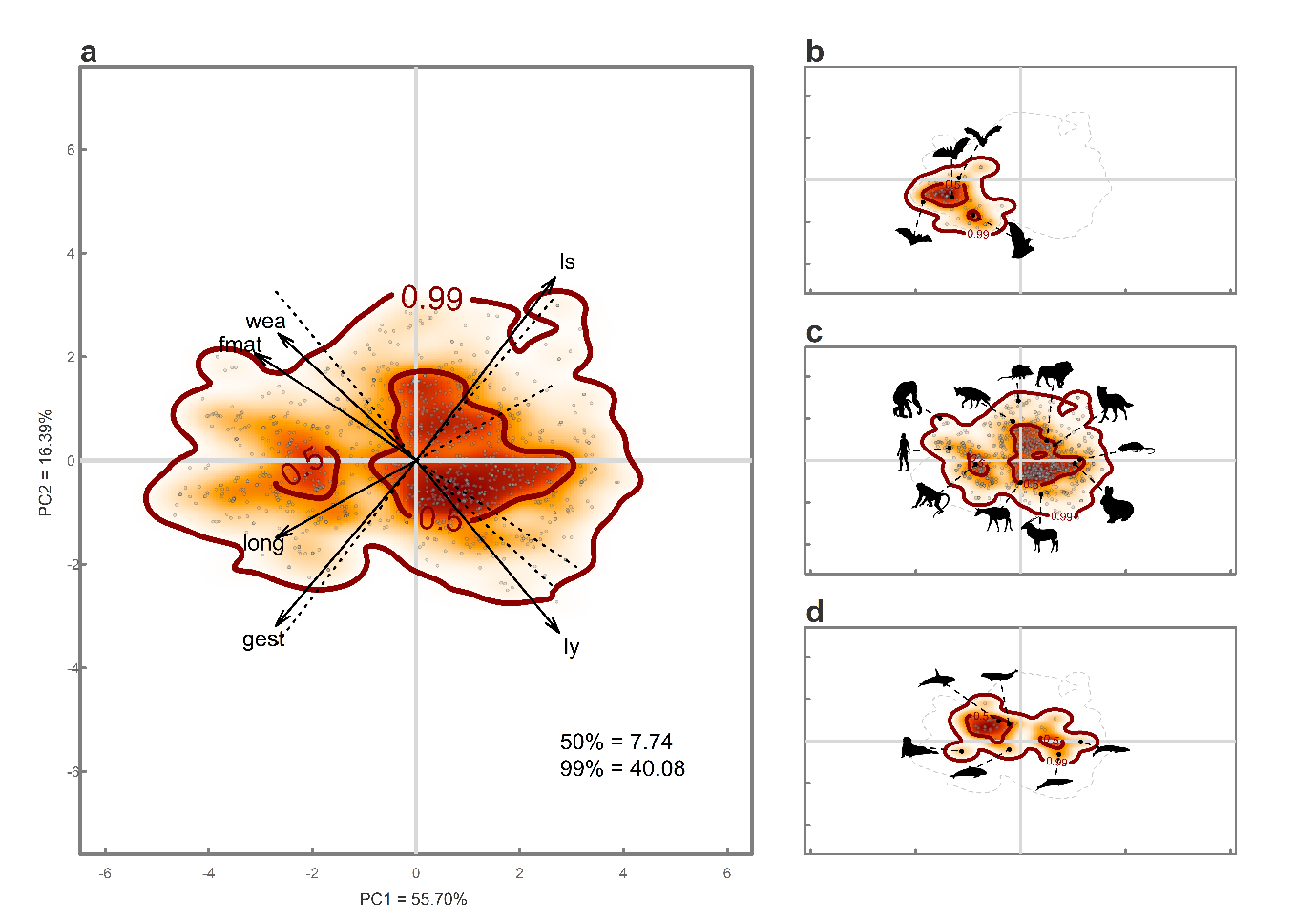
 **Supplementary material Fig. 2 | Mammalian life history space based on complete observation. a,** pattern in space occupation of global mammal in a life history space built considering the complete set of traits (i.e. non imputed traits; N =1,293; see Supplementary methods 2). Arrows indicate the loadings of each trait in the PCA. The legends show the amount of functional space (i.e. functional richness) occupied at 50% and 99% probabilities (indicated by thick contour lines). **b,** pattern in space occupation of aerial mammals in the life history space built on complete traits; **c,** pattern in space occupation of terrestrial mammals in the life history space built on complete traits; **d,** pattern in space occupation of aquatic mammals in the life history space built on complete traits. In all panels, the colour gradient (red, orange, and white) depicts different density of species in the space (red areas have higher density of species). wea, weaning length; fmat, time to reach female maturity; ls, size of litter; gest, gestation length; long, longevity; ly, number of litters yearly. Silhouettes of characteristic species from each realm were downloaded from PhyloPic (http://www.phylopic.org/).

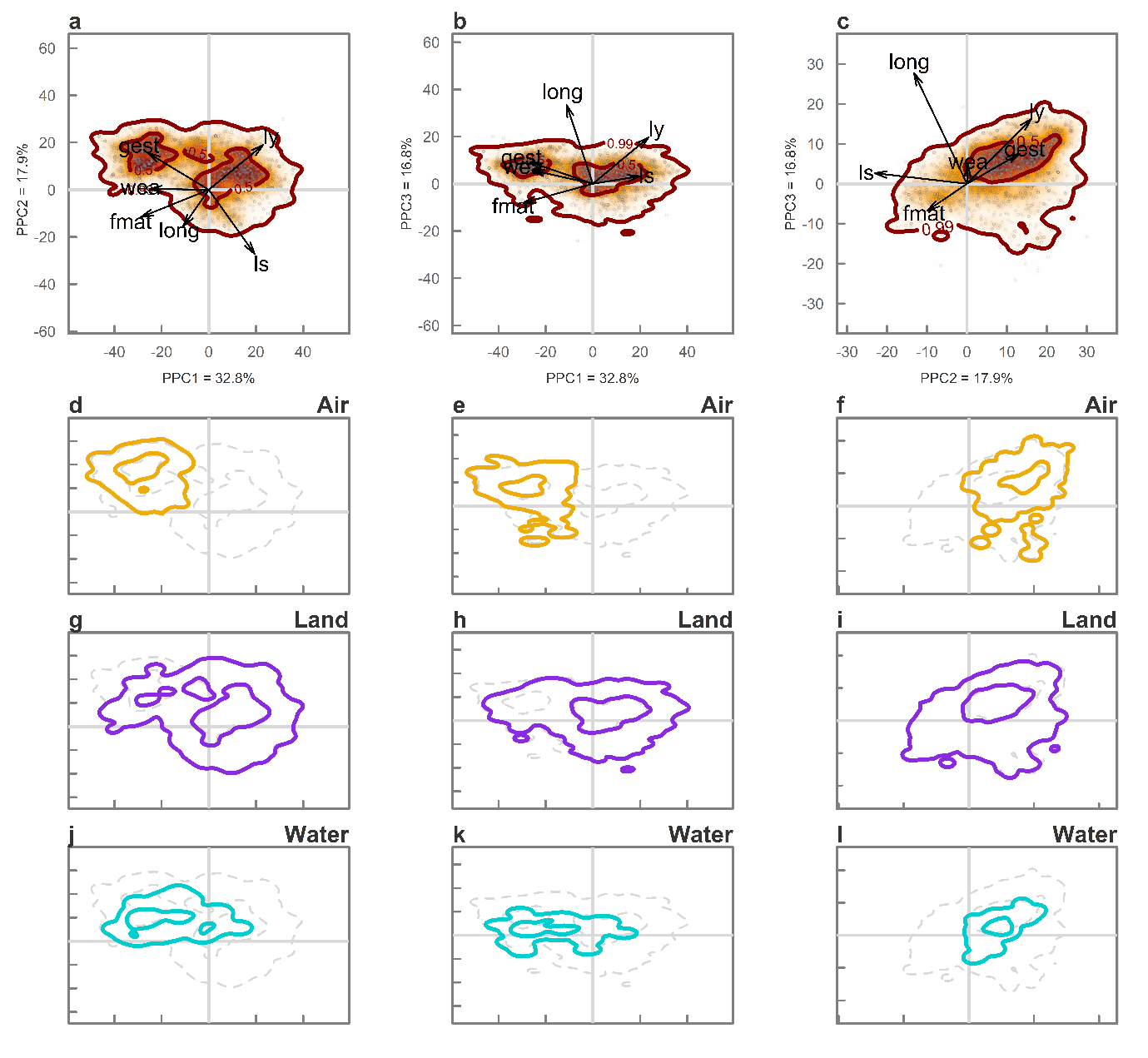
 **Supplementary material Fig. 3 | Mammalian phylogenetic corrected life history space (Phyl-space). a,** pattern in space occupation of global mammal in the first two dimensions (PPC1 and PPC2) of the phylogenetically corrected life history space built using a phylogenetically informed PCA (*i.e.,* Phyl-space; N =3,438; see Supplementary methods 2 and Extended data Table 1). **b,** pattern in space occupation of global mammal considering the first and third dimension (PPC1 and PPC3) of the Phyl-space; **c,** pattern in space occupation of global mammal considering the second and third dimension (PPC1 and PPC3) of the Phyl-space. Arrows indicate the loadings of each trait in the pPCA. The legends show the amount of functional space (*i.e*. functional richness) occupied at 50% and 99% probabilities (indicated by thick contour lines). **d. – g. – j.**, pattern in space occupation of aerial (yellow lines), terrestrial (purple lines), and aquatic (light blue lines) mammals in the first two dimension of the Phyl-space. **e. – h. – k.**, pattern in space occupation of aerial, terrestrial, and aquatic mammals in the first and third dimension of the Phyl-space. **f. – i. – l.**, pattern in space occupation of aerial, terrestrial, and aquatic mammals in the second and third dimension of the Phyl-space. wea, weaning length; fmat, time to reach female maturity; ls, size of litter; gest, gestation length; long, longevity; ly, number of litters yearly.

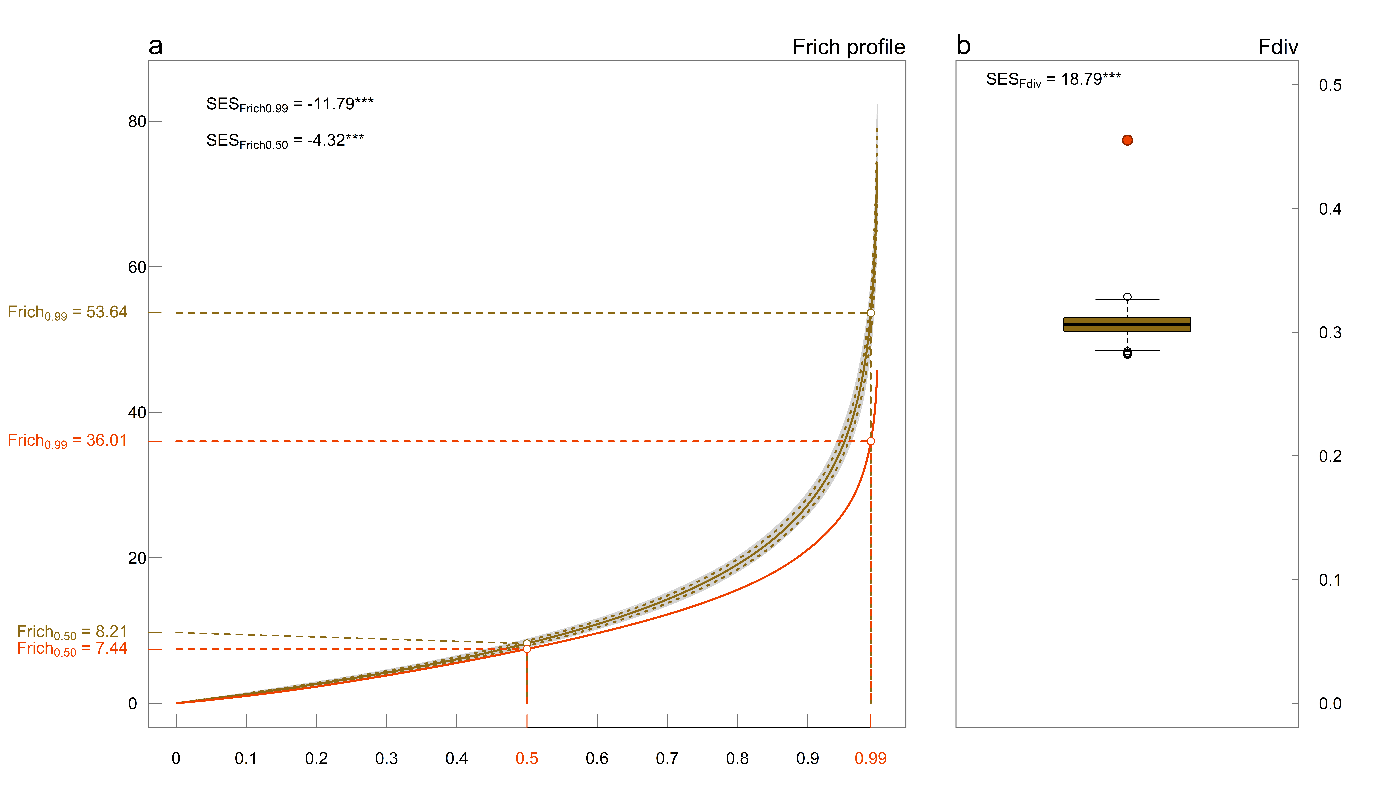
 **Supplementary material Fig. 4** **| C**[**omparison of the occupation of the life history space with multivariate normal distributions**](https://www.nature.com/articles/s41586-021-03871-y/figures/8)**. a,** Functional richness profile (amount of functional space occupied by quantiles of the functional spectra in the mammalian life history space. Gold lines represent the mean, 2.5% and 97.5% quantiles of the functional richness profiles of null models (n = 999) representing multivariate normal distributions with equivalent parameters (means and standard deviations) than the observed data; red line represents the functional richness profile of the observed species. Each repetition of the null model (n = 999) is represented with a thin grey line. The values of functional richness for the 0.5 and 0.99 quantiles of observed and simulated profiles are shown for comparison. Standardize Effect Size (SES) values are reported for both 0.5 and 0.99 quantile which were computed as (observed value – mean (simulated values))/s.d.(simulated values). **b,** functional divergence (representing the degree to which the density of species in the trait space is distributed towards the extremes of the distribution of species in the mammalian life history space. The red point shows the observed functional divergence, and the golden boxplot contains all simulated functional divergence of null models (n = 999). The centre bounds of box, and whiskers of the boxplot represent the median, 25^th^ and 75^th^ percentiles, and 1.5 times the interquartile range***,*** respectively. SES value for functional divergence was computed as above and it is reported in the upper part of the panel. In both panels, two-sided p values were estimated by confronting the value of the SES with the cumulative normal distribution with mean = 0 and standard deviation = 1; *** = p-values <0.001.

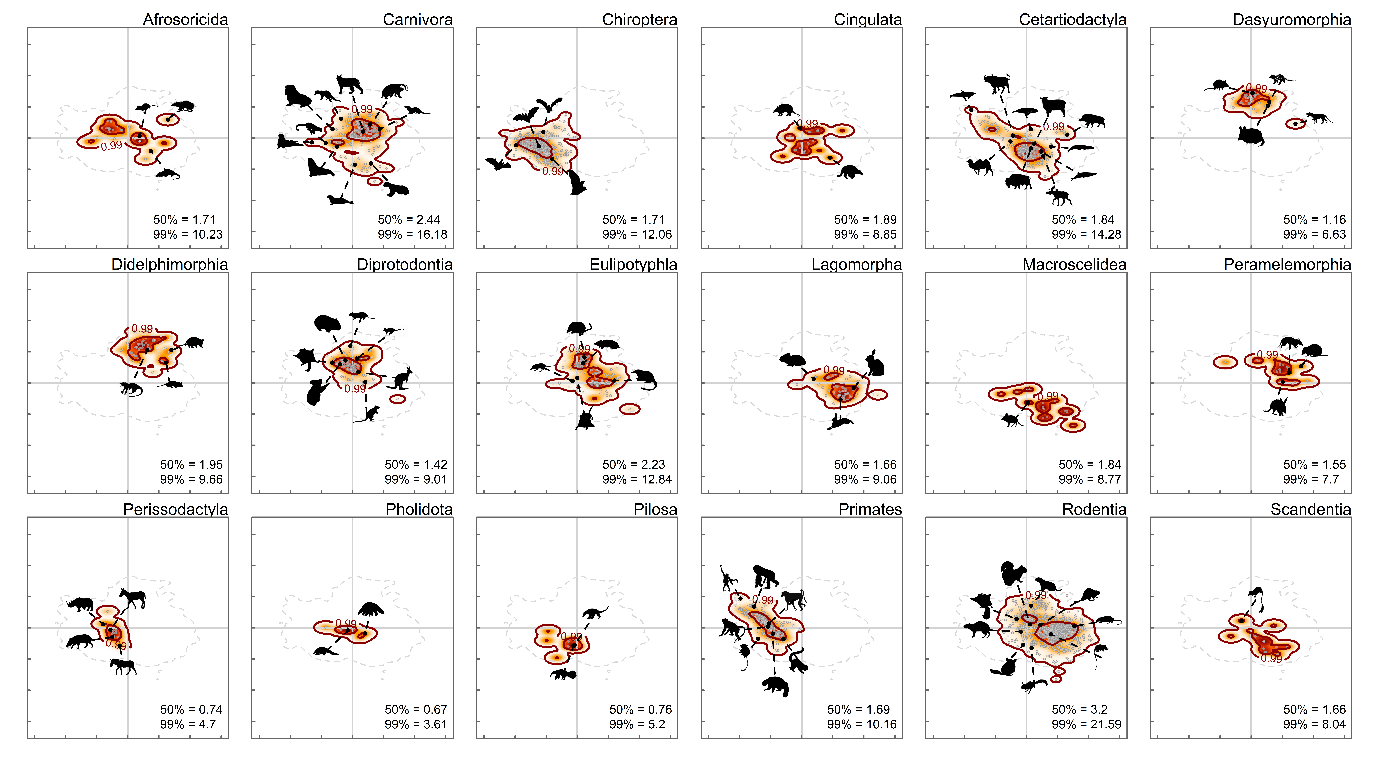

**Supplementary material Fig. 5 | Mammalian orders in life history space.** Pattern in space occupation of mammalian orders (with more than 5 species). Probabilistic species distributions in the spaces defined by the two first principal components (PC1 and PC2) of PCA considering life history traits for all mammals. The colour gradient (red, orange, and white) depicts different density of species in the space (red areas have higher density of species). The legends show the amount of functional space (i.e. functional richness) occupied at 50% and 99% probabilities (indicated by thick contour lines). Silhouettes of characteristic species from each order were downloaded from PhyloPic (http://www.phylopic.org/).

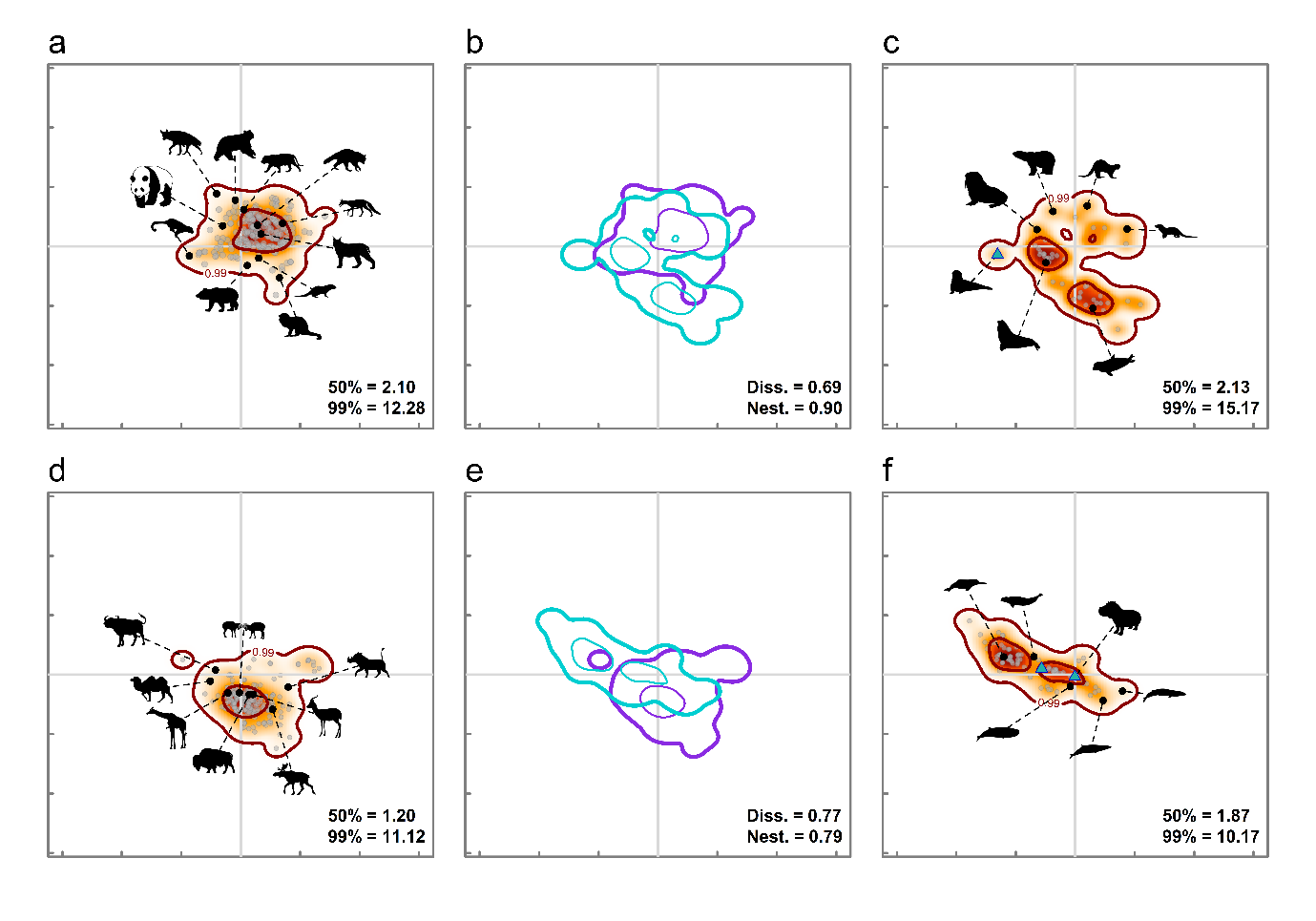

**Supplementary material Fig. 6 | Dissimilarities between orders exploiting two realms. a-c,** Carnivora species patterns: **a,** pattern in space occupation of terrestrial species; **b,** overlap-based dissimilarity between terrestrial (purple) and aquatic (light blue) carnivores; **c,** pattern in space occupation of semi-aquatic species. Carnivora contains only semi-aquatic (i.e., transitional) species, except for the sea otter which is considered as “fully” aquatic (triangle in panel **c**). **d – f,** Cetartiodactyla species: **d,** pattern in space occupation of terrestrial species; **e,** overlap-based dissimilarity between terrestrial (purple) and aquatic (light blue) Cetartiodactyla species; **f,** pattern in space occupation of aquatic species. Cetartiodactyla contains only aquatic species, except for the hippos which are labelled as semi-aquatic species (triangles in panel **f**). **b and e,** dissimilarities are highlighted at 50% (dashed line) and 99% (full line) of total probability, with the legend containing dissimilarity (Diss.) and nestedness (Nest.) values. **a, c, d, f,** in the patterns in space occupation, the colour gradient (red, orange, and white) depicts different density of species in the space (red areas have higher density of species). The legends show the amount of functional space (i.e. functional richness) occupied at 50% and 99% probabilities (indicated by thick contour lines). Silhouettes of characteristic species from each realm were downloaded from PhyloPic (http://www.phylopic.org/).

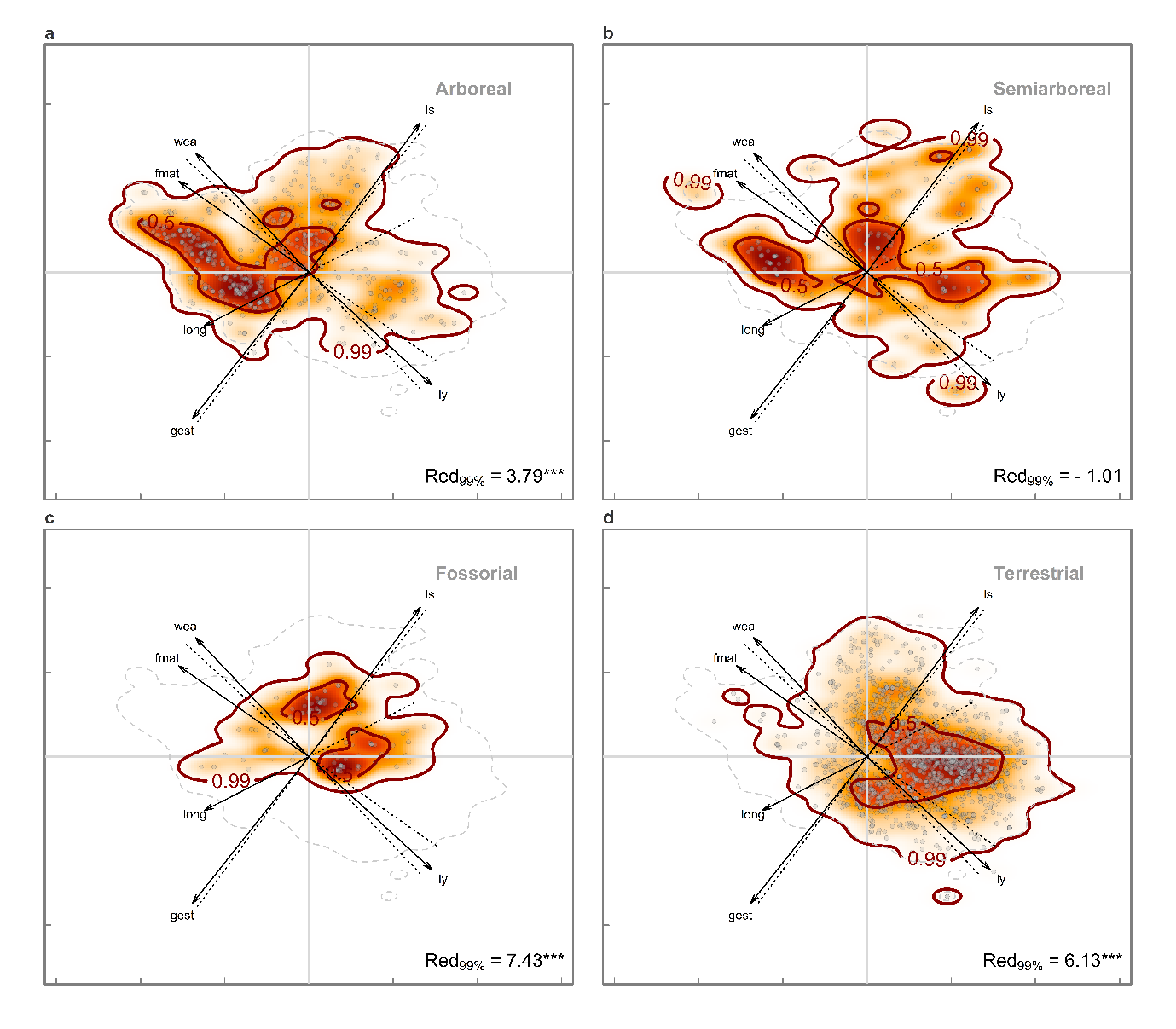

**Supplementary material Fig. 7 | Terrestrial mode-of-life show distinct life history structures.** Patterns in space occupation of arboreal (**a**), semiarboreal (**b**), fossorial (**c**), and species confined on land, or ‘fully’ terrestrial (**d**) mammals. Thick contours indicate 50% and 99% probabilities. Arrows indicate the loadings of each trait in the PCA. Legends show functional redundancy of the mode-of-life calculated as absolute values of the Standardized Effect Size (Red_99%_). SES was calculated considering the amount of functional space occupied by observed and randomized realm at 99% probability. * is SES p-value < 0.05; ** SES p-value < 0.01; *** SES p-value < 0.001. In all panels, the colour gradient (red, orange, and white) depicts different density of species in the space (red areas have higher density of species). wea, weaning length; fmat, time to reach female maturity; ls, size of litter; gest, gestation length; long, longevity; ly, number of litters yearly.

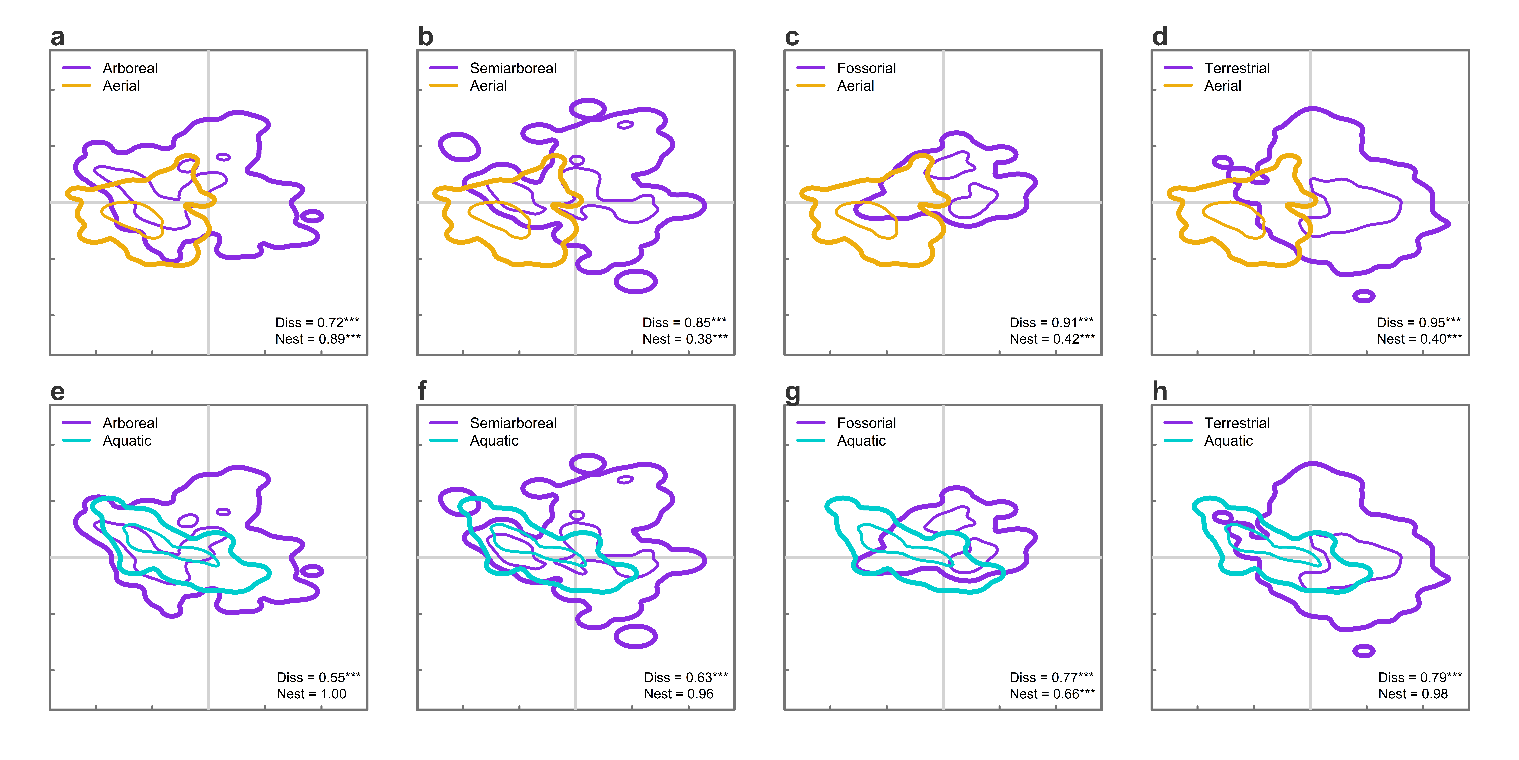

**Supplementary material Fig. 8 | Arboreal, semiarboreal, and fossorial mode-of-life show strategies similar to aerial and aquatic realms.** Overlap-based dissimilarities between terrestrial (purple), aerial (light orange), and aquatic (light blue) species distributions in the life history space discriminating the different mode-of-life present in the terrestrial realm. Specifically: **a,e,** dissimilarity of arboreal species with aerial and aquatic realms; **b,f,** dissimilarity of semiarboreal species with aerial and aquatic realms; **c,g,** dissimilarity of fossorial species with aerial and aquatic realms; **d,h,** dissimilarity of species confined on land (terrestrial) with aerial and aquatic realms. Each life history structure is highlighted at 50% (fine lines) and 99% (thick lines) of total probability. SES of dissimilarity and nestedness values for each pairwise comparisons are reported in the legend together with SES p-values (* SES p-value < 0.05; ** SES p-value < 0.01; *** SES p-value < 0.001).

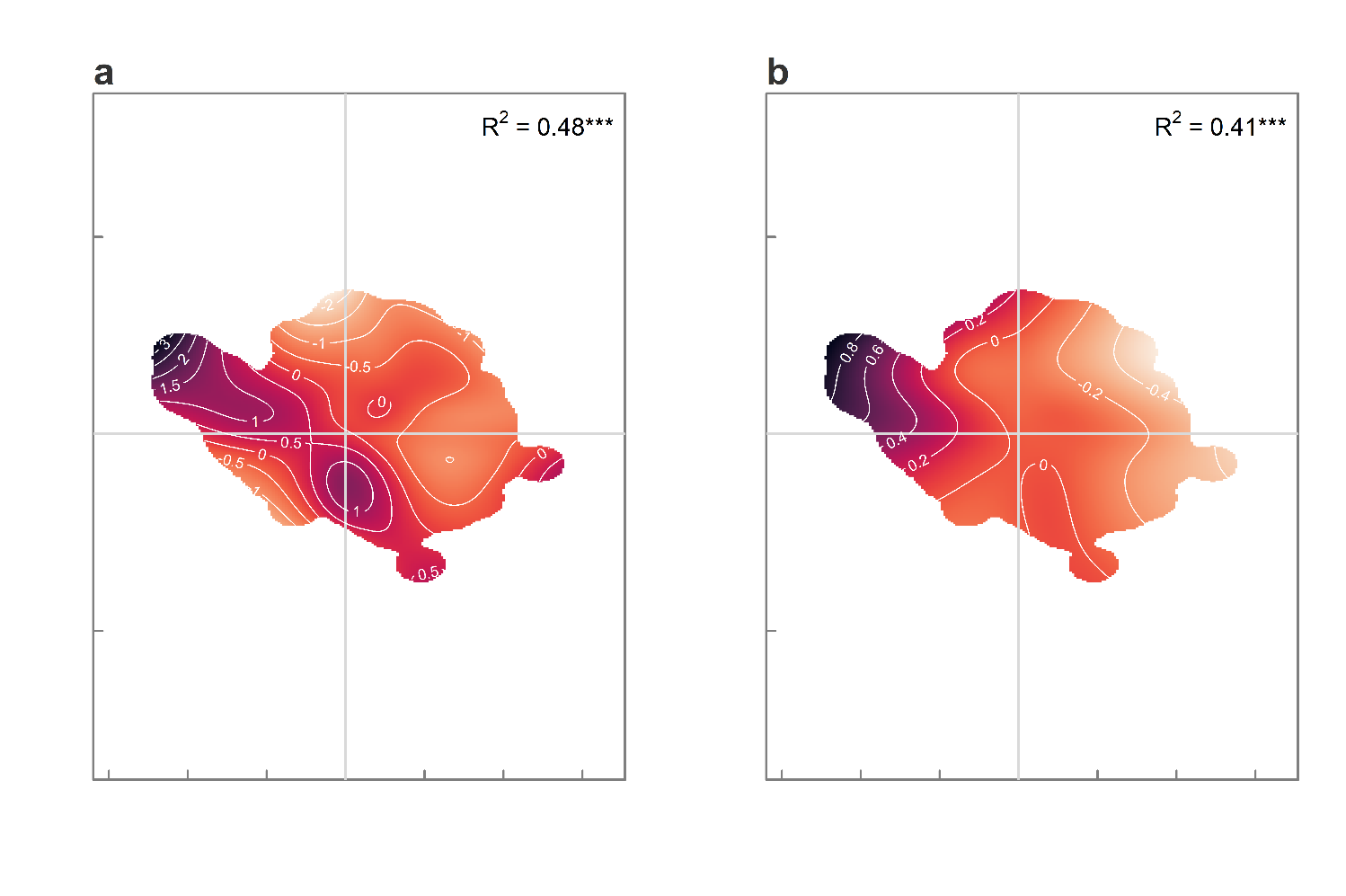

**Supplementary material Fig. 9 | Brain mass of terrestrial species as a function of species position in the life history space. a.,** Observed brain mass distribution in the life history space; **b.**, brain – body mass residuals distribution in the life history space (Supplementary methods 4). Observed brain mass and brain-body mass residuals were used as a response variable. The position of terrestrial species in the space was used as a predictor in both generalized additive models (GAMs). The adjusted r-squared of each model is reported in the upper part of the panels; *** p-value of model predictors < 0.001. Darker tones represent bigger brain mass (**a.**) or higher brain-body mass residuals (**b.**), and viceversa. White contour lines represent the thresholds of different brain mass or (**a.**) or brain-body mass residuals (**b.**).

**Supplementary material Table 1 | Principal component analyses (PCAs) of global mammalian life history traits.** Proportion of variance (P. Variance), eigenvalues, loadings of principal components, and angles (degrees) between the eigenvectors of all pairs of traits in the selected principal components in four different PCAs. **a.**, PCA performed on body mass–corrected residuals computed from the imputed set of life history for the 3,438 species considered in the main text (expressed graphically in the main text [Fig.](https://www.nature.com/articles/nature16489#Fig2) 1). **b. - d.**, PCAs used to test the methodological choices made during the construction of the life history space as described in Supplementary methods 2. **b.**, PCA performed on body mass–corrected residuals computed only for the spaces with complete records (1,293 species) for the six traits used in the main analysis (Expressed graphically in Supplementary Fig. 6). **c.**, PCA performed on the same set of imputed life history traits without body mass correction is the PCA carried out on the same set of imputed traits used in the main analysis without body mass correction (3,438 species). **d.**, phylogenetic PCA (Phyl-PCA) performed on the same body mass–corrected residuals used for the main analysis (3,438 species).

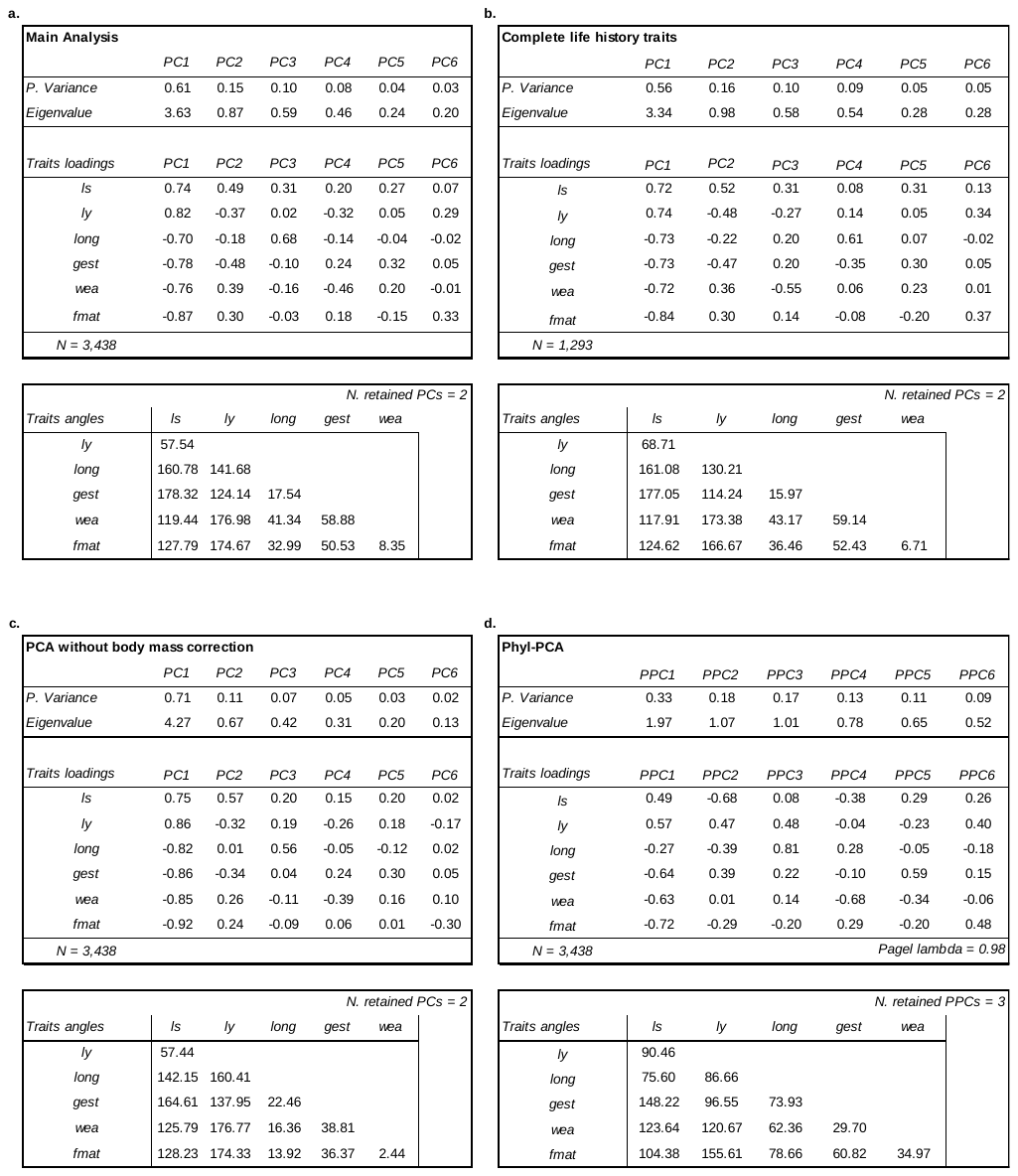

**Supplementary material Table 2** | **Functional redundancy and dissimilarity patterns within environmental realms. a.**, Functional redundancy is calculated as the standardized effect size (SES) of observed functional richness (F.richness) and functional richness computed randomizing species assignment to the realm. **b.**, Pairwise dissimilarities (Dissim.) and nestedness values (Nest.) between realms. *Air* is the aerial realm, *Water* is the aquatic realm, *Land* is the terrestrial realm, and *SA* are the semi-aquatic species transitioning between realms. In all tables, observed value (Obs), mean of the randomized values of 999 null models (Rand. mean), standard deviation of the randomized values of the of 999 null models (Rand. Sd.), standardized effect size (SES), and p-value of the standardized effect size (p-value SES).

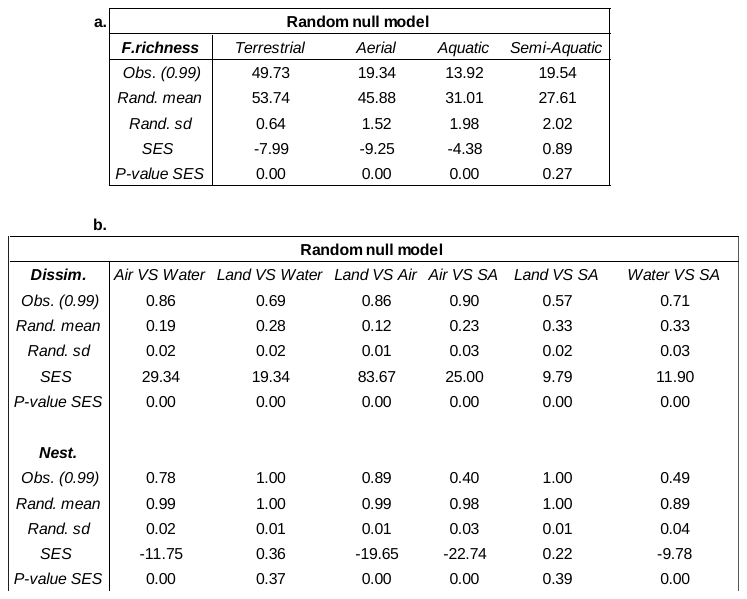

**Supplementary material Table 3 | Functional redundancy and dissimilarity patterns within terrestrial realms and across arboreal, semiarboreal and fossorial mode-of-life. a.**, Functional redundancy is calculated as the standardized effect size (SES) of observed functional richness (F.richness) and functional richness computed randomizing species assignment to different mode-of-life. **b.**, Pairwise dissimilarities (Dissim.) and nestedness values (Nest.) between realms and mode-of-life. *Arb.* is the arboreal mode-of-life*, Semiarb.* is semiarboreal mode-of-life, *Foss.* is fossorial mode-of-life, *Land* are species confined on land, *Air* is the aerial realm, *Water* is the aquatic realm, and *SA* are the semi-aquatic species. In all tables, observed value (Obs), mean of the randomized values of 999 null models (Rand. mean), standard deviation of the randomized values of the of 999 null models (Rand. Sd.), standardized effect size (SES), and p-value of the standardized effect size (p-value SES).

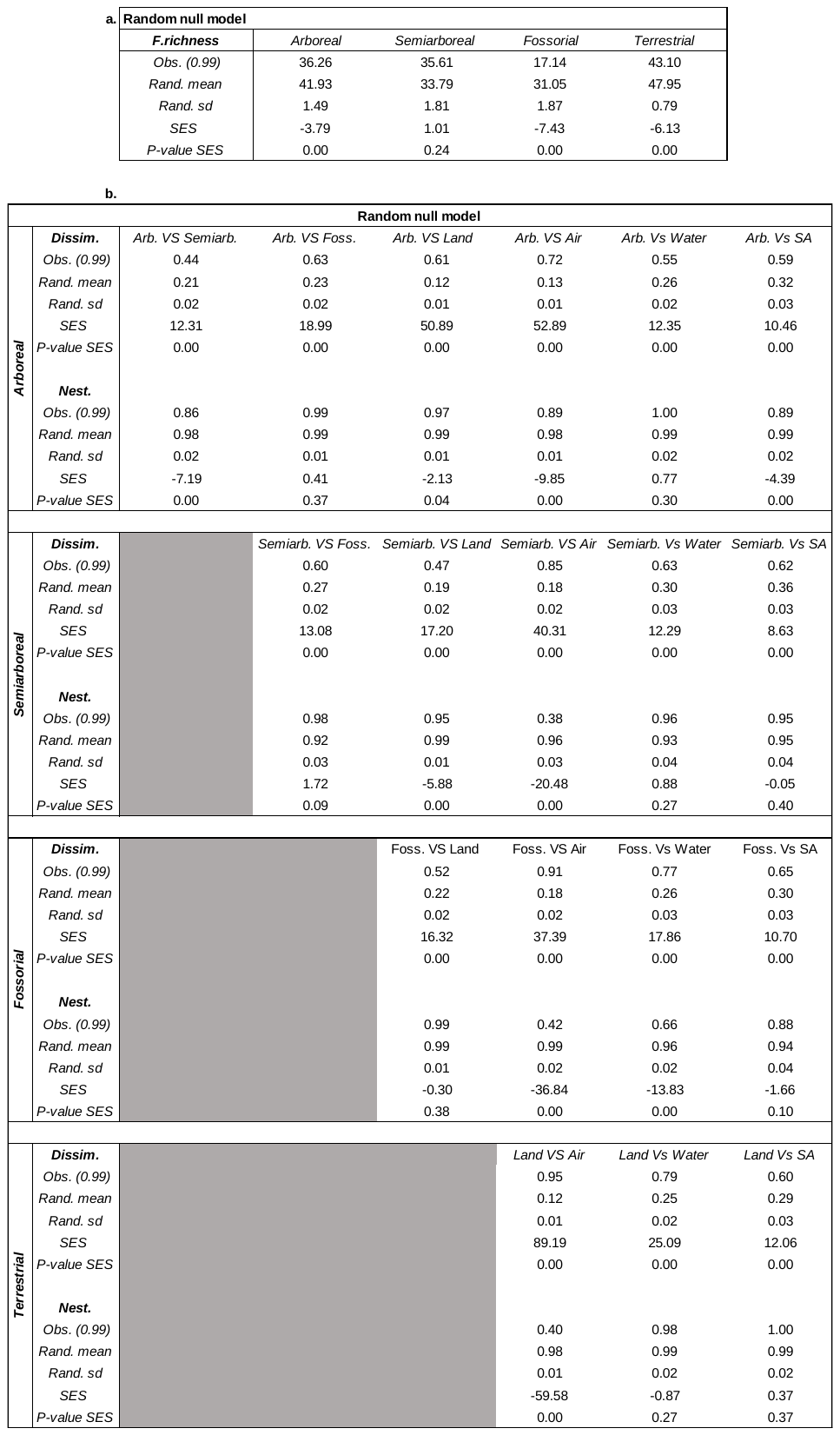
 **Supplementary material Table 4 | Statistical significance of differences among species confined on land and arboreal, semiarboreal, and fossorial species**. Model outputs of the linear mixed effect models using mean score of species in each order as response variables, mode-of-life as fixed effect, and mammalian orders as random intercept. The model was performed to test for statistically significant differences between species confined on land and species with the considered mode-of-life along the two dimensions of the life history space. For each model the reference category were orders with species confined on land.

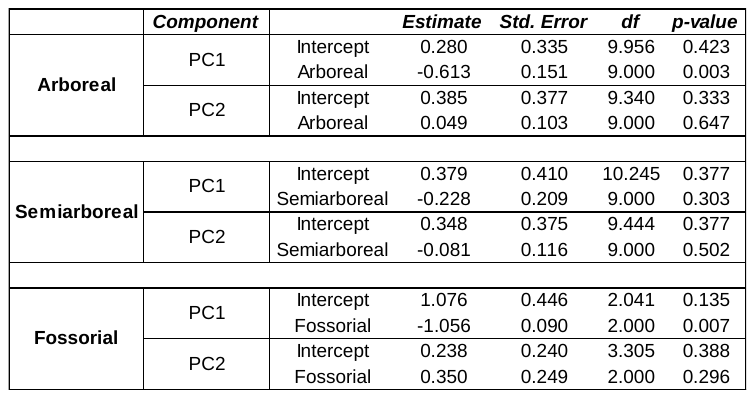
