## Supplementary methods for "Global Diversity in Mammalian Life Histories: Environmental Realms and Evolutionary Adaptations"

**Supplementary methods 1: Reliability of the imputation procedure**

To estimate the reliability of the position of species with imputed trait values in the trait space, we explored the performance of the imputation procedure both considering individual traits and species positioning in the life history space. We used the same procedure already performed by Carmona *et al.* (2021a): we tested imputation performance by artificially removing trait values from the 1,293 species with complete trait measurements. Within this subset of species with complete information, we reproduced a pattern of missing values consistent with the one in the original dataset. For that, we randomly extracted 10 species with incomplete information and superimposed their patterns of missing trait values to 10 randomly selected species from the subset of species with complete empirical information. In doing so, we built a matrix identical to the one used to impute the original data (N = 3,438), with the only exception of the 10 randomly selected species with complete empirical information to which we added artificial missing values. Using this matrix, we performed the same phylogenetically informed imputation procedure used to impute the original dataset as described in the Methods section of the main text (See “Data collection and trait imputation” subsection). Then, we used the artificial set of imputed life histories to account for body mass using the same procedure described in the Methods section of the main text. Specifically, we extracted intercept and slope for each ordinary linear regression performed in the original analyses and used it to regress single artificial life history traits and observed species adult body mass. It is important to stress that mean body mass records were empirically measured for all species used in the dataset. Then, we extracted artificial body mass–corrected residuals, and we centred it using the same mean and standard deviation of original body mass–corrected residuals. Additionally, we used artificial body mass–corrected residuals to build an artificial life history space as done for the original dataset (see main text Methods section, subsection “Construction of the global life-history traits space”). We estimated the performance of the imputation on species position in the life history space using the normalized root mean square error (NRMSE), which expresses the average distance between original and artificial positions of species on each axis of the life history space as a proportion of the range of the axis (Penone *et al.* 2014). Specifically, we compared the position of the 10 randomly selected species in the original life history space without removing data (real position of the species) and after artificial removal and imputation of trait information. We repeated this procedure 9.100 times, to achieve 70 repetitions for each of the species with complete empirical trait information and attained an estimation of the average NRMSE value across repetitions and for each dimension of the life history space.

Our results show that even in case of traits missing for some species, the imputation procedure allows to estimate the position of the species in the life history space with high accuracy. The NRMSE for each dimension is:

- Component 1: 0.028 (0.003 st.dev.)
- Component 2: 0.017 (0.001 st.dev.)

Consequently, on average, the imputation procedure retrieves the real position of a species with an error of around 2.8% of the range for the first component and 1.7% for the second component.

**Supplementary methods 2: Reliability of the methodological choices in the life-history space**

**Life history space based on complete records.** To further test the reliability of our life history space based on imputed data, we compared the original life history space with one built considering only complete species records (*i.e.,* not imputed records, N = 1,293). We extracted the set of species with complete traits observation and performed the same procedure used to build the original life history space (see Main text, Methods section and “Construction of the global life history traits space.” subsection). Specifically, we accounted for the effect of body mass by performing a series of linear regressions between single complete life history traits and species adult body mass. Then, we used the body mass–corrected residuals to construct a life history space based only on complete species records by performing principal component analysis (PCA, Supplementary material Fig. 2; and Supplementary material Table 1).

Then, we compared the life history space built using only complete records (N = 1,293) with the original life history space built using imputed data (N = 3,438). We extracted the position of the species with common information from the original life history space and we performed a Procrustes rotation using the *protest* function of the *‘vegan’* R package and considering 999 permutations (Oksanen *et al.* 2020). High values in the Procrustes rotation would indicate that the relative position of species is similar between the space built using only species with a complete set of observation and the original life history space built using imputed data. Additionally, we tested whether traits correlation in the two spaces were consistent by estimating the angles between the loadings of all pairs of traits in the original life history space and in the life history space with complete records (see Bueno *et al.* 2023 and Carmona *et al.* 2021a; Supplementary material Table 1). We then estimated the correlation between these angles, with high correlations indicating that the inferred relationships between pairs are similar for both spaces.

Procrustes analysis showed significant correlation (r = 0.98; p < 0.001) suggesting a very high correspondence between the imputed and complete life history spaces and demonstrating that species positioned in the space is not influenced by the imputation procedure. The correlation between loadings’ angles in the two spaces was almost perfect and statistically significant (r = 0.998, p < 0.001), demonstrating that imputation did not influence the inferred relationships between life history traits (Supplementary material Fig. 2; Supplementary material Table 1).

**Body mass correction in the life history space.** We further tested the reliability of our life history space by testing whether accounting for body mass influenced species position in the life history space. In doing so, we compared the original life history space (which was built using body mass–corrected residuals, see Main text, Methods section, “Construction of the global life-history traits space.” subsection) to a space built using the same set of imputed life history traits (N = 3,438) but without body mass correction. We computed the life history space without body mass correction (hereafter Bm-Space) by performing a PCA directly on scaled and centred life history traits (Supplementary material Table 1). Then, we compared the so built Bm-Space with the original life history space following the same procedure described above.

Procrustes analysis showed significant correlation (r = 0.79; p < 0.001) suggesting a very high correspondence between Bm- and life history spaces. The correlation between loadings’ angles in Bm-Space and original life history space was high, positive, and statistically significant (r = 0.98, p < 0.001). These results, suggest that original life history covariation and species positioning in life history space is not strongly influenced by body mass correction procedure.

**Phylogeny in the life history space.** Considering that both environmental filtering and evolutionary history influence trait covariation(Blomberg & Garland Jr 2002; Southwood 1988; Stearns 1992), similar evolutionary histories among species can constrain the life history trait values among taxa. In doing so, evolutionary history can influence how species occupy the life history space, producing patterns that strongly depend on phylogeny rather than environmental realm conditions. Therefore, to account for evolutionary differences between species, we computed a phylogenetically corrected space (*i.e.,* Phyl-space) using the same size-corrected life history traits we used to build the observed life history space. To compute the Phyl-space we performed a phylogenetically informed PCA (PPCA, Revell 2009). PPCA considers phylogenetic non-independence of species by inversely weighting the PCA covariance matrix estimated from the phylogeny, thus removing the expected phylogenetic correlation among traits (Polly *et al.* 2013; Revell 2009). Differently from PCA, PPCA does not produce major axes of trait variation, but rather major axes of phylogenetically-independent trait variation centred on the basal node of the phylogeny (Polly *et al.* 2013; Revell 2009). Species scores are produced through a rigid rotation of the original trait data in the PPCA space; therefore, scores conserve original trait variance. Consequently, PPCA axes can be correlated although their phylogenetic correlations will be zero (Polly *et al.* 2013; Revell 2009). PPCA was performed considering the same phylogeny described in the main text (section Methods, subsection “Data collection and trait imputation”). We used the *phyl.pca* function of the *‘phytools’* R package (Revell 2012). To define the Phyl-space, we extracted the first three PPCs which together retained 67.46% of the total life history variance (Supplementary material Table 1 and Supplementary material Fig. 3).

Then, we compared the Phyl-space with the life history space built without performing the phylogenetic correction. First, we performed a Procrustes rotation as mentioned in the previous section. Furthermore, we estimated the correlation between dissimilarities in the position among pairs of species in the life history space with dissimilarities in the Phyl-space. We estimated species pairwise dissimilarity on each space by computing Euclidean distances between PCs and PPCs axes, and we tested how these similarities were correlated by performing a Mantel test using the *‘vegan’* R package (Oksanen *et al.* 2020). High values in the Procrustes rotation as well as high correlations between the two dissimilarity matrices would indicate that the relative position of species is similar in the two compared spaces. We tested whether traits correlation in the two spaces were consistent. For this, we estimated the angles between the loadings of all pairs of traits in the original life history space and in the Phyl-space (Supplementary material Table 1) and estimated the correlation between these angles. High correlations between the angles estimated in both spaces would indicate that the inferred relationships between pairs are similar for both spaces.

Additionally, we tested for the correlation between phylogenetic and trait dissimilarities between pairs of species. We extracted species with complete set of life history raw traits (non-imputed, N = 1,293 species) and we pruned the phylogenetic tree to match species subset using the *drop.tip* function of *‘phyltools’* R package. We computed pairwise distances between pairs of tips in the phylogenetic tree using the *cophenetic.phylo* function of the R package *‘ape’*(Paradis & Schliep 2019). Then, we computed dissimilarities between raw life history traits using Euclidean distance in the *vegdist* function of the R package *‘vegan’* (Oksanen *et al.* 2020). Then, we tested for the correlation between species phylogenetic and functional dissimilarities by performing a Mantel test. We performed the same procedure considering phylogenetic and life history traits dissimilarities considering the full set of imputed plus body-mass corrected life history traits used to build the life history space.

Procrustes analysis showed significant correlation (r = 0.95; p < 0.001) suggesting a very high correspondence between the Phyl-space and the life history spaces. Consistently, dissimilarities between species’ positions in the life history space and Phyl-space showed a strong correlation (ρ = 0.94; Mantel test: p < 0.001), suggesting that phylogenetic history is not the main driver of species positioning in the life history space. The correlation between loadings’ angles in the two spaces was statistically significant (r = 0.85, p < 0.001), demonstrating that phylogeny doesn’t influence relationships between life history traits shaping the main dimensions of life history variations (Supplementary material Fig. 3).

Finally, the phylogenetic and functional dissimilarities between species showed a low correlation for both non-imputed (ρ = 0.15; Mantel test: p < 0.001; N = 1,293) and imputed with body mass correction (ρ = 0.25; Mantel test: p < 0.001; N = 3,438) life history traits, suggesting that dissimilarity in phylogenetic history of species is not strongly driving species dissimilarity in life history strategies. Altogether, these results demonstrate a strong correspondence between life history space and Phyl-space, suggesting the lack of strong evolutionary influence in shaping species positioning in the life history space.

**Supplementary methods 3: Aquatic dependency Index (ADI)**

Different databases relay on different interpretation of degree of dependency to water needed to define a species as aquatic. For instance, according to Worms database (WoRMS Editorial Board 2020), seal species emulates dolphins in their strict aquatic classification. On the other hand, species like rhesus macaque (*Macaca mulatta)* that occasionally have been observed swimming in water, are considered semi-aquatic. Inversely, IUCN (IUCN 2017) classifies the same *Macaca mulatta* species, as fully terrestrial, whereas seals species are considered as semi-aquatic, and dolphins are fully aquatic. Aiming to overcome these incongruences in the classification of fully aquatic and semi-aquatic species, we summarised species dependency to water by creating the Aquatic Dependency index (AD Index). AD Index broadly summarises whether a species depends for important portion of their day (*i.e.,* feeding, hide in den) and life (*e.g.,* reproduction) in water. The index answer to 5 five binary questions: *(i) Does the species copulates in water?* *(ii) Does the species give birth in water?*; *(iii) Does the species wean the offspring in water?*; *(iv) Does the species mandatorily search for food in water?*; *(v) Does the species find refuge in water (i.e. doesn’t have a den on land)?* Answers to these questions were found throughout a literature review(Hood 2020; Perrin *et al.* 2009; University of Michigan 2022). Positive answers were summed to produce an index ranging from 0 to a maximum degree of aquatic dependency of 5. In case the answer was not clear or not present, we answered using the available information of the closest taxa to that species (*e.g.,* other representatives of the same family or genus). We classified all species scoring between 1 and 4 in the ADIndex as semi-aquatic and all species scoring 5 as fully aquatic (*i.e.,* aquatic realm).

| **ADI** | **Level 1** | **Level 2** | **Level 3** | **Level 4** | **Level 5** |
| --- | --- | --- | --- | --- | --- |
| Number of species (% on the total of aquatic and semi-aquatic)  N = 143 | 17 (11.9%) | 24 (16.8%) | 16 (11.2%) | 1 (0.7%) | 85 (59.4%) |
| Classification | SEMI-Aquatic | SEMI-Aquatic | SEMI-Aquatic | SEMI-Aquatic | Aquatic |
| Orders | Afrosoricida;  Carnivora;  Eulipotyphla;  Rodentia. | Carnivora;  Monotremata;  Rodentia. | Carnivora;  Cetartiodactyla. | Carnivora. | Carnivora;  Cetartiodactyla;  Sirenia. |

**Supplementary methods 4: Mapping species encephalization in the life history space**

We explored whether high encephalization (Barton & Capellini 2011; González-Lagos *et al.* 2010; Pérez‐Barbería *et al.* 2007; Reader & Laland 2002) allows to exploit portion of the life history space otherwise not successful for the terrestrial realm. For this we downloaded adult brain mass from the Combine imputed database (Soria *et al.* 2021). We extracted brain mass information for the subset of 1,054 species with complete records. Then, we accounted for the covariation between brain mass and body dimension by extracting the residuals from an ordinary linear regression between log-transformed brain and body mass (*i.e.,* independent variable) of terrestrial mammalian species. We mapped observed log-transformed brain mass and brain-mass residuals in response to species position in the life history space following Carmona *et al.*, (2021b). For this, we used two generalized additive models (GAM), where observed brain mass and brain-mass residuals and brain mass was used as response variable, respectively. In both models, species coordinates in the life history space were used as explanatory variables and by computing 6 basis functions (k) of thin plate regression splines basis type, considering both main effect of each PCs composing the life history space as well as the interaction between them. As smoothing parameter estimation, we used the REML method. The GAM was computed using the r package ‘mgcv’ (Wood 2017). Model output considering brain mass as response variable was the following:

| *Parametric coefficients:* | | | |
| --- | --- | --- | --- |
|  | Estimate | Std. Error | p-value |
| Intercept | 2.474 | 0.051 | 0.000 |
| *Approximate significance of smooth terms:* | | |  |
|  | edf | p-value |  |
| PC1 | 3.995 | 0.000 |  |
| PC2 | 4.508 | 0.000 |  |
| PC1:PC2 | 13.898 | 0.000 |  |
| R-sq.(adj) = 0.483 | | |  |

Model output considering brain-body mass residuals as response variable was the following:

|  | Parametric coefficients: | | | |
| --- | --- | --- | --- | --- |
|  | | Estimate | Std. Error | p-value |
| Intercept | | -0.025 | 0.013 | 0.065 |
| *Approximate significance of smooth terms:* | | | |  |
|  | | edf | p-value |  |
| PC1 | | 4.141 | 0.000 |  |
| PC2 | | 4.476 | 0.000 |  |
| PC1:PC2 | | 8.236 | 0.006 |  |
| R-sq.(adj) = 0.410 | | | |  |

These results show that terrestrial species with higher brain mass cluster in the slower portion of the life history space (Supplementary material Fig. 9). Results are confirmed after accounting for allometric covariation between brain and body sizes, suggesting that results are independent from body dimensions (Barton & Capellini 2011). Models demonstrated that species positions in the life history space is significantly related to species encephalization.

**Supplementary methods references**

Barton, R.A. & Capellini, I. (2011). Maternal investment, life histories, and the costs of brain growth in mammals. *Proc. Natl. Acad. Sci. U.S.A.*, 108, 6169–6174.

Blomberg, S.P. & Garland Jr, T. (2002). Tempo and mode in evolution: phylogenetic inertia, adaptation and comparative methods. *J. Evol. Biol.*, 15, 899–910.

Bueno, C.G., Toussaint, A., Träger, S., Díaz, S., Moora, M., Munson, A.D., *et al.* (2023). Reply to: The importance of trait selection in ecology. *Nature*, 618, E31–E34.

Carmona, C.P., Bueno, C.G., Toussaint, A., Träger, S., Díaz, S., Moora, M., *et al.* (2021a). Fine-root traits in the global spectrum of plant form and function. *Nature*, 597, 683–687.

Carmona, C.P., Tamme, R., Pärtel, M., Bello, F. de, Brosse, S., Capdevila, P., *et al.* (2021b). Erosion of global functional diversity across the tree of life. *Sci. Adv.*

González-Lagos, C., Sol, D. & Reader, S.M. (2010). Large-brained mammals live longer. *j. Evol. Biol.*, 23, 1064–1074.

Hood, G.A. (2020). *Semi-aquatic Mammals: Ecology and Biology*. JHU Press.

IUCN. (2017). *The IUCN Red List of Threatened Species. Version 3*. Available at: https://www.iucnredlist.org. Last accessed 23 September 2020.

Oksanen, J., Blanchet, F.G., Friendly, M., Kindt, R., Legendre, P., McGlinn, D., *et al.* (2020). *vegan: Community Ecology Package*. Available at: https://CRAN.R-project.org/package=vegan. Last accessed .

Paradis, E. & Schliep, K. (2019). ape 5.0: an environment for modern phylogenetics and evolutionary analyses in R. *Bioinformatics*, 35, 526–528.

Penone, C., Davidson, A.D., Shoemaker, K.T., Di Marco, M., Rondinini, C., Brooks, T.M., *et al.* (2014). Imputation of missing data in life-history trait datasets: Which approach performs the best? *Methods Ecol. Evol.*, 5, 961–970.

Pérez‐Barbería, F.J., Shultz, S. & Dunbar, R.I.M. (2007). Evidence for coevolution of sociality and relative brain size in three orders of mammals. *Evolution*, 61, 2811–2821.

Perrin, W.F., Würsig, B. & Thewissen, J.G.M. (2009). *Encyclopedia of marine mammals*. Elsevier Inc.

Polly, P.D., Lawing, A.M., Fabre, A.-C. & Goswami, A. (2013). Phylogenetic Principal Components Analysis and Geometric Morphometrics. *Hystrix*, 24.

Reader, S.M. & Laland, K.N. (2002). Social intelligence, innovation, and enhanced brain size in primates. *Proc. Natl. Acad. Sci. U.S.A.*, 99, 4436–4441.

Revell, L.J. (2009). Size-Correction and Principal Components for Interspecific Comparative Studies. *Evolution*, 63, 3258–3268.

Revell, L.J. (2012). phytools: an R package for phylogenetic comparative biology (and other things). *Methods Ecol. Evol.*, 3, 217–223.

Soria, C.D., Pacifici, M., Di Marco, M., Stephen, S.M. & Rondinini, C. (2021). COMBINE: a coalesced mammal database of intrinsic and extrinsic traits. *Ecology*, 102, e03344.

Southwood, T.R.E. (1988). Tactics, Strategies and Templets. *Oikos*, 52, 3–18.

Stearns, S.C. (1992). *The evolution of life histories*. Oxford university press Oxford.

University of Michigan. (2022). *Animal Diversity Web (ADW)*. Available at: https://animaldiversity.org. Last accessed .

Wood, S.N. (2017). *Generalized Additive Models: An Introduction with R, Second Edition*. CRC Press.

WoRMS Editorial Board. (2020). *World Register of Marine Species*. Available at: https://www.marinespecies.org. Last accessed 1 September 2020.
